# Finding the rhythm: Humans exploit nonlinear intrinsic dynamics of compliant systems in periodic interaction tasks

**DOI:** 10.1101/2023.08.31.555654

**Authors:** Annika Schmidt, Marion Forano, Arne Sachtler, Davide Calzolari, David Franklin, Alin Albu-Schäffer

## Abstract

Everyday activities, like jumping on a trampoline or using a swing-stick, show that humans seemingly effortless support systems in their intrinsically preferred motions. Although this observation seems obvious, data-based evidence proving that humans indeed match system dynamics has been lacking, since everyday objects usually exhibit complex, nonlinear dynamics, which are in general not analytically solvable. Recent insights in the field of nonlinear mode theory and the development of a tool to compute modes for nonlinear systems enabled us to investigate human strategies to excite periodic motions in the interaction with nonlinear systems. In the setup of a high score game, participants interacted with differently configured virtual compliant double pendulum systems through a haptic joystick. Through the joystick, the user could command positions to a motor link connected to the pendulum by a spring and received resulting spring forces in return to convey the feeling of holding a flexible stick. The participants were asked to alternately hit two targets located on the computed nonlinear mode of the system as often as possible. All participants intuitively exploited the elasticity of the system by choosing a *holding strategy* of the motor link and only compensate for energy losses with small motions. In this way, the intrinsic dynamics of the double pendulum system were exploited leading to the predicted fast motions along the nonlinear modes. The human strategy stayed consistent when decreasing the target size or increasing the mass of the lower pendulum link, i.e., changing the dynamics. Consequently, the presented research provides data-based evidence that humans can indeed estimate the nonlinear dynamics of system and intuitively exploit these. Additionally, the introduction to nonlinear modes and ways to compute them could be a powerful tool for further investigations on human capabilities and strategies in periodic interactions with nonlinear systems.

**Author summary:** Without thinking about it, humans interact with a wide variety of objects in everyday life. This includes objects with very complex nonlinear dynamics such as flexible rods or ropes. Since it is not trivial to enforce trajectories far away from the system’s intrinsic motions and frequencies, it is likely that humans explore and, whenever possible, exploit the natural dynamics of the system. By using a tool to predict the trajectories of systems with nonlinear dynamics, we collected human user data to validate this hypothesis for repetitive tasks with a virtual double pendulum. Indeed our research showed that humans supported mechanical systems in their respective intrinsic motions and were able to intuitively match the systems’ eigenfrequencies. In doing so, only little control effort and motion was needed from the users, which could aids to save energy and mental resources. Since both these aspects are limiting factors in continuous tasks, it seems to have an evolutionary benefit that humans are very capable in estimating and exploiting the natural dynamics of compliant systems and tune their own control strategy to be be synchronized to the controlled system.

## Introduction

Humans are very dexterous and versatile in using their upper limbs in everyday tasks. Arm trajectories are usually carried out without conscious planning, while the muscle force and stiffness are flexibly adjusted to accommodate different tasks. Moreover, humans can handle countless objects with ease, even when these objects are complex in shape, flexible in material, have multiple degrees of freedom (DOFs), and highly nonlinear dynamic behavior. Seemingly effortless humans can move around and manipulate or interact with objects, like jumping on a trampoline, exciting a swing stick or hammering a nail. For all these motions and interactions, it is most likely that both the dynamics of the human arm [1] as well as the dynamics of an interacted object [2] influence the human strategy how arm motions are carried out and tools are used [3, 4]. Studies suggest that in interactions with objects with internal DOFs, humans not only have an internal model of their own body dynamics but likewise form an internal model of the object they interact with [5]. This implies that humans must be able to estimate the dynamics of external objects through haptic interactions. This idea is also supported by recent works from the research groups of Hogan and Sternad [1, 2, 6–8]. One of their studies showed that humans simplified constrained motion tasks by taking advantage of interactive dynamics and tuning their hand impedance to the system rather than applying precise force control [6]. Further, a cup balancing task experiment showed that participants always chose similar starting positions to initialize the task [7], indicating that the participants could get an intuitive feeling of the system dynamics. Even in experiments with a system with very complex dynamics, such as a whip, humans might excite the simple intrinsic modes of the system to hit a target [8]. Thus, for interactions with dynamic systems, experiments support the idea that humans can effectively estimate and make use of the system dynamics.

However, analyzing how and to what extent humans really match or even exploit object dynamics is very challenging. The human sensorimotor system is extremely complex, starting with noisy sensors, considerable information delays, and the need to coordinate multiple muscles, all accumulating to a highly nonlinear system that is very difficult, if not impossible, to model in its entirety. Even the much simpler, purely mechanical objects that humans interact with in everyday life usually exhibit complex, often nonlinear dynamics. This makes it hard to predict the intrinsic behavior of an object, such that it is impossible to say to which extent the human might make use of it. Nevertheless, observations and research show that humans are capable of exciting oscillatory motions like bouncing a ball supporting the intrinsic system dynamics to reach stability [9–11]. The used internal physical models are most likely intuitively learned based on Newtonian principles [12, 13]. Robotic experiments showed that an optimal control model considering the nonlinear dynamics of the human arm could predict human performance in such interactions realistically, which could not be accounted for by a linear model [14]. This emphasizes the importance of accounting for nonlinearities in dynamic interactions, not only of the human but also of the interacted objects. Furthermore, the work of Sternad and colleagues showed that in dynamic interactions, humans are indeed sensitive to object dynamics and favor strategies that make interactions stable and predictable [2]. Motions that are inherently easy to stabilize are the intrinsic motions of a system, i.e., the motions that a system is inclined to do based on its physical properties. Considering only periodic intrinsic motions, a well-known subclass of such are the eigenmodes of a system. Eigenmodes have been well studied for the linear case [15, 16], but usually everyday examples do not adhere to such idealized cases. However, recent advances in the field of nonlinear eigenmodes [17, 18] have shown that even nonlinear systems such as the double pendulum, the textbook example for chaotic behavior [19], in fact exhibit nonlinear modes. As part of these advances, our group has developed a numerical *mode tool* that is able to compute these nonlinear modes for compliant systems, i.e., the motions a system would intrinsically sustain in the conservative (frictionless) case [20, 21]. This enables us to predict the preferred motion, even of nonlinear systems, and allows us to analyze if and how well humans match and exploit the inherent system dynamics.

We hypothesize that in dynamic rhythmic interactions, humans will intuitively favor strategies that exploit the intrinsic system motions, i.e., the modes of a system or object. In line with past findings that humans seek stability and predictability in dynamic interactions [2], such control strategies seem likely as supporting intrinsic dynamics make it easier to stabilize motions [22] and thus require less interventions and, in turn, less human effort, decreasing the load on the human sensorimotor control.

We directly test the hypothesis that humans can effectively estimate the dynamics of a compliant system and will tune their control strategy to match and support the dynamics accordingly. As the system for the dynamic interaction with the human, we chose a compliant double pendulum for the following reasons: 1) A compliant double pendulum is the most basic simplification of a flexible object, such as a whip or rope, and could be extended in future work to incorporate more links for interactions with more complex flexible objects. 2) The compliant double pendulum also captures essential dynamic properties of a human limb moving in a vertical plane, e.g., as first approximation modeling the human’s upper and lower arm by a double pendulum with some potential energy due to gravity and elasticity. In this sense, studying how humans can estimate, excite, and exploit intrinsic motions of a double pendulum is interesting from two perspectives: as a complex, possibly chaotic interaction object and as a simplified but very relevant model of human biomechanics. Thus, we designed a human user study in which participants interacted with a virtual compliant double pendulum through a haptic joystick as interface. Following a high-score game setup, the participants were asked to excite the pendulum to hit two targets aligned with the intrinsically preferred motion of the pendulum system as often as possible in a given time. The experiment results showed that all users intuitively supported the intrinsic motions of the system by tuning their control strategy to be in sync with the system eigenfrequency. These findings stayed consistent for changed system dynamics and increased task difficulty. Even when the required task was not aligned with the intrinsic system motion path, participants made use of the compliance in the system. These observations provide new data-based evidence to support the intuitive knowledge that humans very effectively estimate and exploit the dynamics of compliant systems.

## Results

Before the human user experiments, the double pendulum system was analyzed with the developed *mode tool* [17, 20, 21] to derive the expected trajectory of the system for the conservative case, i.e., the intrinsic motion the system will follow in the frictionless case. Additionally, in simulation, the behavior of the double pendulum systems was characterized through a frequency sweep and random excitation. Based on the system characterization and initial pilot experiment, we identified baseline strategies that we expect participants in the human user study could use to achieve the task of hitting between two targets with the endpoint of the virtual pendulum (Fig. 1). The initial experiment (Experiment 1) focused on identifying and characterizing the human user’s control approach by comparing the excited pendulum motions to the theoretical mode and the defined baseline strategies. Following, Experiment 2 investigated the robustness of the human control approach by altering the mass of the lower pendulum link, i.e., the system dynamics or increasing the task difficulty through smaller target sizes. Each experiment variation, i.e., changed mass or target size, was tested with one half of the participants. Finally, Experiment 3 tested if and how the human control strategy would change if the locations of the targets to hit were not aligned with the trajectory the double pendulum system would intrinsically follow.

**Fig 1.**
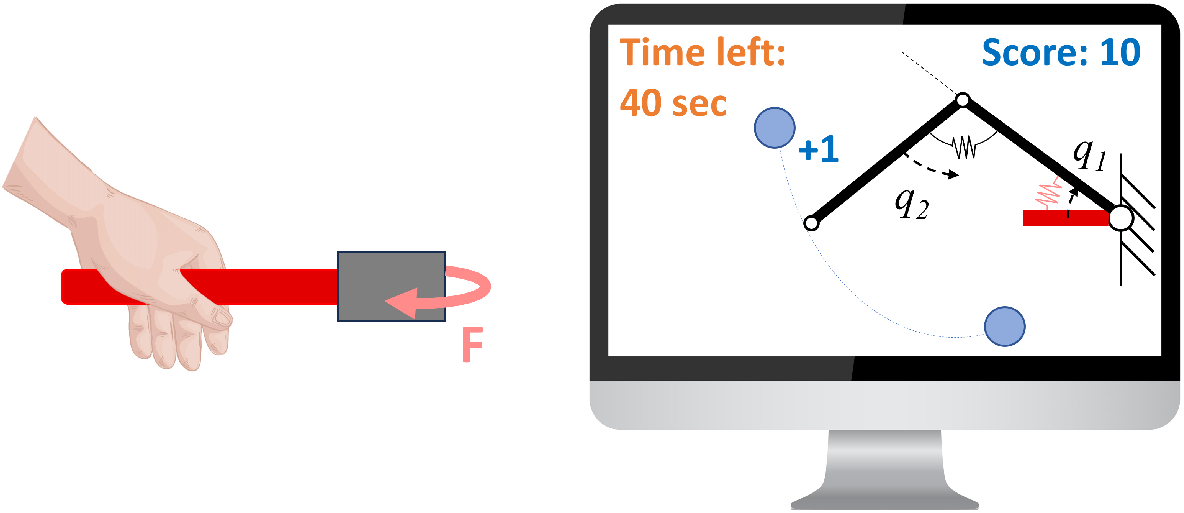
Illustration of the experimental setup: The participant controls a virtual motor link with a 1:1 position mapping through a haptic joystick (red). The motor link is coupled to a virtual compliant double pendulum by a spring. The occurring forces *F* of that spring are reflected to the user as force feedback. The goal is to hit two targets (blue circles) with the lower link endpoint as often as possible within a given time of 40 s.

### Nonlinear modes of the conservative double pendulum

As the theory of nonlinear normal modes is crucial for this work and the user study’s design, we briefly introduce the topic, but the full theory and details are covered within [17, 18].

Starting from a well-known example case for linear oscillations, consider the simple system of a pendulum swinging horizontally in a plane, i.e., without gravity’s influence. It has the *mass m*_1_ located in the middle of the link and a rotational, linear spring with stiffness *k*_1_ in its joint (Fig. 2a). When the mass is deflected and then released, assuming the conservative frictionless case, the system will naturally oscillate with a constant frequency that is mainly determined by the mass and stiffness of the system. This is the system’s so-called eigenfrequency *ω*, and the resulting oscillation is the eigenmode or normal mode of this pendulum [15, 16]. In joint space, this mode can be described by an eigenvector, which would be, for the given simple case, just a straight line that increases in length for greater deflections, i.e., higher energy levels (Fig. 2d). Adding a second link with mass *m*_2_ to the simple pendulum system leads to a double pendulum (Fig 2b,c), which is usually the textbook example of a nonlinear system with chaotic behavior. However, our theory and numerical tools for nonlinear modes show that if we release the conservative double pendulum from certain initial angles with zero velocity, it will robustly follow a well-defined periodic trajectory without deviation and without exhibiting chaotic behavior [17, 18]. In particular, it will perform rest-to-rest motions, returning to the initial rest point after a full cycle (Fig. 2e, green). In this nonlinear case, the eigenvector turns into a curve. The values of the needed initial conditions change for different energy levels, but there will always be a condition that leads to this modal motion. The collection of initial values that lead to this one kind of intrinsic mode is called *Generator* as they generate the periodic motions for all energy levels (Fig. 2, blue). The union of all trajectories of the Generator is called *Eigenmanifold*. In simple terms, the mode is the intrinsic regular motion that a system will follow in the conservative case if initialized from the adequate values, i.e., from the generator.

**Fig 2.**
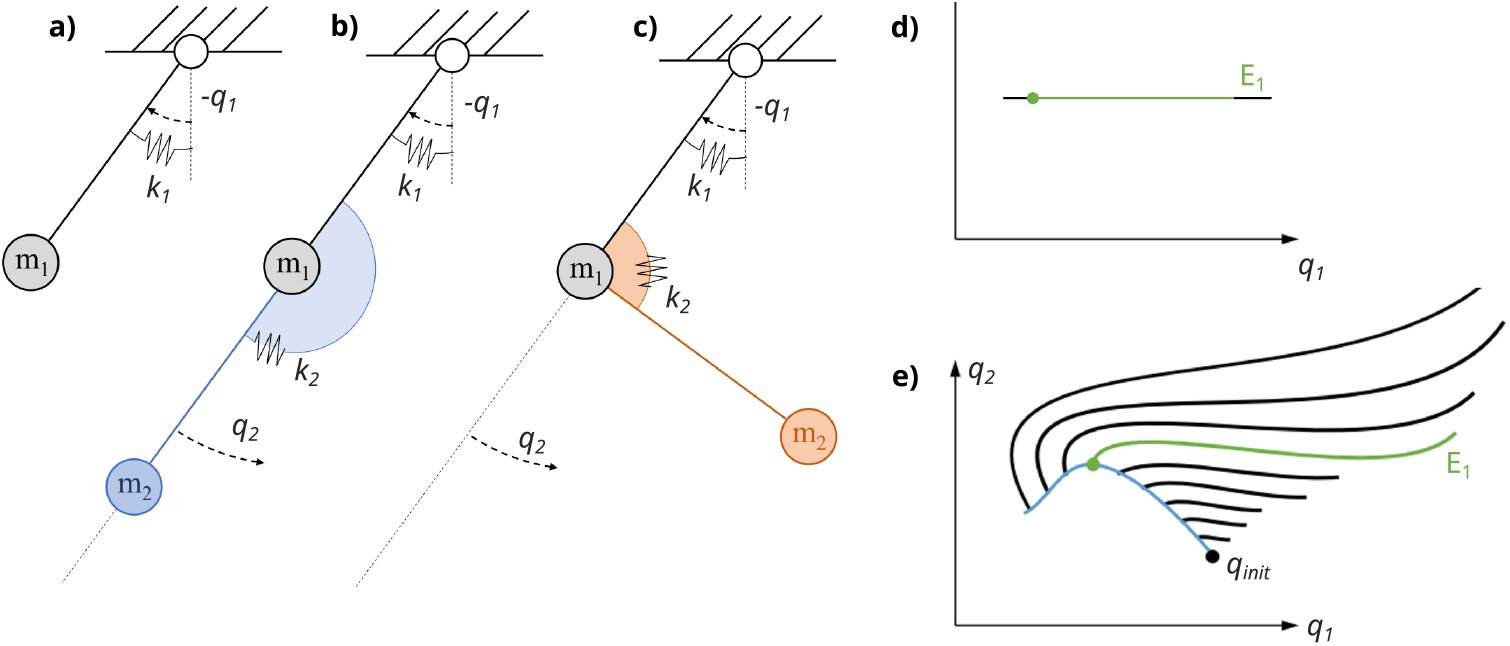
**a)** Schematic simple pendulum with the link-centered mass *m*_1_ and a rotational springs with stiffness *k*_1_ in its origin. The link angle *q*_1_ is defined with respect to the origin. Adding a second link with mass *m*_2_ through another spring of stiffness *k*_2_ leads to a double pendulum, where the spring rest position is either set to **b)** 0° or **c)** 90°. The lower link angle *q*_2_ is defined relative to the upper link. **d)** Schematic illustration of the linear mode for a 1-DOF system, i.e., the simple pendulum with coordinates *q*_1_. With increasing energy the mode increases linearly, where *E*_1_ (green) indicates the trajectory for one defined energy level. **e)** Expanding the system to 2-DOF with a nonlinear mode illustrates that the shape of a nonlinear mode changes with energy. Nevertheless, for every energy level, an initial condition can be found that leads to a mode trajectory. The collection of these initial values *q*_init_ is called *Generator* (blue) and one mode for an arbitrary energy level *E*_1_ is indicated in green. The union of all trajectories of the Generator is called *Eigenmanifold* (image adapted from [17]).

For a more realistic case, where the system encounters friction and other energy losses, a controller needs to add energy to keep the system on the mode by injecting the energy lost in each cycle. This is analogous to a swing on the playground that needs to be pushed at the endpoint to keep swinging with constant amplitude. Multiple robotic experiments [20, 23, 24] have shown that moving a system along its modes is very energy efficient, as the system is supported in a motion it is intrinsically inclined to follow and therefore only little control effort is needed to sustain this motion. Inspired by these findings and the observations of Hogan and Sternad in various works that humans seek predictability and low sensorimotor effort in dynamic interactions [1, 2, 6–8], we hypothesize that humans intuitively choose a control strategy to drive a compliant system that matches the intrinsic system dynamics as these are easier to stabilize. To test this hypothesis, we need to compare the motion that the human excites in a system during a dynamic interaction with the intrinsic motion, i.e. a nonlinear normal mode, that the system would exhibit in the conservative case. Since nonlinear modes cannot be derived analytically for nonlinear systems, we use the newly developed *mode tool* to numerically compute the nonlinear modes of compliant systems [20, 21]. As detailed in the Introduction, the already introduced double pendulum with rotational springs in each joint is chosen as an example case for the human user study (Fig. 2a).

To not limit the investigation of the human strategy to the interaction with only one single system, two different configurations of the compliant double pendulum are considered by varying the equilibrium position of the rotational spring connecting the first and the second link. Measured from the origin of the second link coordinate *q*_2_ as shown in Figure 2a, the first configuration has a spring rest position of 0° (Fig. 2, blue), i.e., fully extended. For the second configuration, the rest position of the spring is 90 ° (Fig. 2, orange), corresponding to a partially flexed arm. In the following, the two configurations will be denoted to as *P0* and *P90*, respectively. The length and mass values of the double pendulum system were roughly based on the dimensions of a human arm. The specific parameter values are detailed in the Method section in Table 2.

For each of the considered pendulum configurations, first, the corresponding nonlinear modes need to be identified as reference to compare the pendulum motion driven by the human in the following experiment (Fig. 3). According to the stated hypothesis, we expect the human to excite each pendulum configuration along its respective mode to exploit the intrinsic system motion to lead to a stable trajectory with minimized effort. Using the *mode tool*, two modes could be identified for each pendulum configuration. In this work, only the first, more stable, mode will be investigated. The computed generators of the first mode for *P0* and *P90* are shown in Figure 3a and 3b in blue, respectively. Since the mode changes its shape with energy, for both configurations, a fixed energy level of 2.5 J was set for the investigation in the human user study. For this energy level, the computed period time of the investigated mode was 1.29 s and 1.08 s for *P0* and *P90*, respectively. This translates to an eigenfrequency of the nonlinear mode at 2.5 J of *f*_res(P0)_ = 0.78 Hz and *f*_res(P90)_ = 0.92 Hz. The resulting mode in joint space is shown in green in Figure 3a and 3b. As visualized, the mode of the *P0* configuration is still relatively close to a linear solution. In contrast, the *P90* mode is clearly nonlinear, so two clearly different cases are considered with the two configurations. The corresponding paths of the double pendulum for the *P0* and *P90* configuration in Cartesian space are shown in Figure 3c and 3d, respectively.

**Fig 3.**
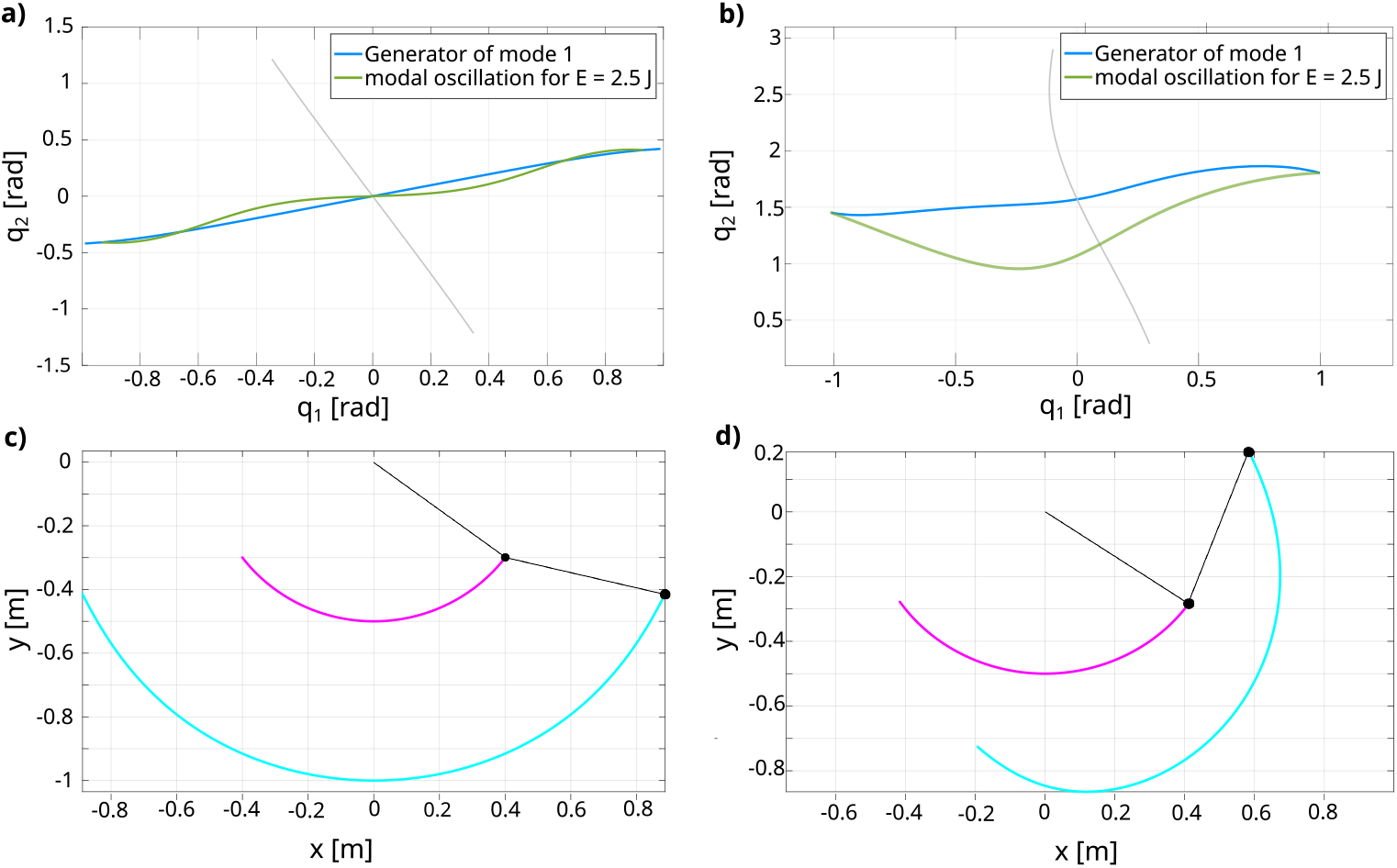
Identified modes for the double pendulum configuration with a **a)** 0° offset (*P0*) and a **b)** 90° offset (*P90*) in the spring connecting the upper and the lower link. The blue line indicates the mode generator for each system, while the green line shows the mode trajectory for an energy level of 2.5 J, which will be used as a reference in the human user study. **c**,**d)** Cartesian trajectories of the link endpoints for the modes of the respective systems *P0* and *P90*.

### Baseline strategies for system excitation

Based on an initial characterization of the double pendulum systems through a frequency sweep (detailed in the method section) and the hypothesis that humans will exploit the intrinsic motions of systems, three *Baseline Strategies* (*BS1-3*) were defined to outline possible control strategies that the human could apply to drive a continuous pendulum motion to move between two targets.

#### BS1: Resonance

The sweep measurements for the system characterization detailed in the method section showed that the biggest system response was triggered with a sine wave input close to the eigenfrequency. For this frequency, the commanded motor link motion was comparatively small to the high amplitudes of the joint trajectory, close to the expected nonlinear mode. In line with our hypothesis, this suggests that humans might intuitively choose the natural frequency of the double pendulum systems to drive their motions. Thus, the first baseline strategy assumes the sine wave (4) with

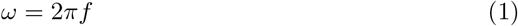

with *f* being the eigenfrequencies *f*_res(*P*0)_ and *f*_res(*P*90)_ for the *P0* and *P90* configuration, respectively, as computed for the conservative case. The amplitude was tuned empirically to *A* = 0.12 such that both targets are hit.

#### BS2: Position Control

A more conservative strategy had been observed in initial pilot experiments, where users first interacted with the system. Here, the motor link was still moved in a sinusoidal-like fashion but with a much slower frequency, such that the elasticity of the springs was not exploited and rather a position control of the upper link was applied. To mimic this behavior in simulation as second baseline for comparison, the frequency of the sine wave from (4) was set to 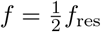 and the amplitude once more tuned empirically to hit the targets.

#### BS3: Bang-bang

In previous research, a similar experiment as presented here had suggested that a bang-bang control law could represent the human behavior [25], which showed to work robustly and optimally efficient in robotic systems [23, 24, 26]. Thus, this control strategy is likewise considered as a possible excitation approach of the human and applied in simulation to obtain a third baseline for comparison. A jump in the motor link *θ* is applied whenever a fixed torque threshold *ϵ*_*τ*_ is crossed, where the torque *τ* arose from the forces of the spring between the motor and the upper pendulum link:

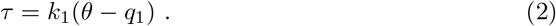

The constants for the threshold and the desired position command were tuned empirically to hit the targets leading to *ϵ*_*τ*_ = 1 and 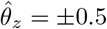 Depending on the sign, the motor link coordinate *θ* is then commanded to do a jump to a desired position 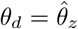according to

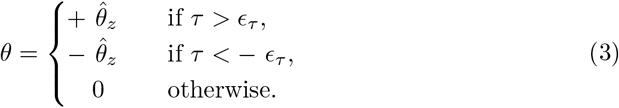

The three presented baseline strategies are considered as possible control approaches to fulfill the defined experiment task of exciting the pendulum systems to repetitively hit between two targets. It needs to be mentioned that these baseline strategies to drive the pendulum motion for the experiment task are idealized solutions, while the human will most likely apply a more flexible distorted motion. Nevertheless, these baselines are intended to provide a way to identify the underlying strategy concept applied by the human by comparing the human data to the ones obtained in the baseline simulations.

### Experiment 1

#### Comparison with expected baseline strategies

To identify the underlying control strategy applied by the participants, the pendulum motions excited through the participants in the user study were compared to the previously defined baseline strategies (Fig. 4). To quantify how well each strategy made use of the intrinsic dynamics of the double pendulum systems, two metrics were especially important: 1) the excited oscillation frequency of the system and 2) the distance of the excited pendulum motion to the expected nonlinear mode of the conservative system set as reference. The congruence was quantified by a *mode metric*, which is detailed in the Method section.

**Fig 4.**
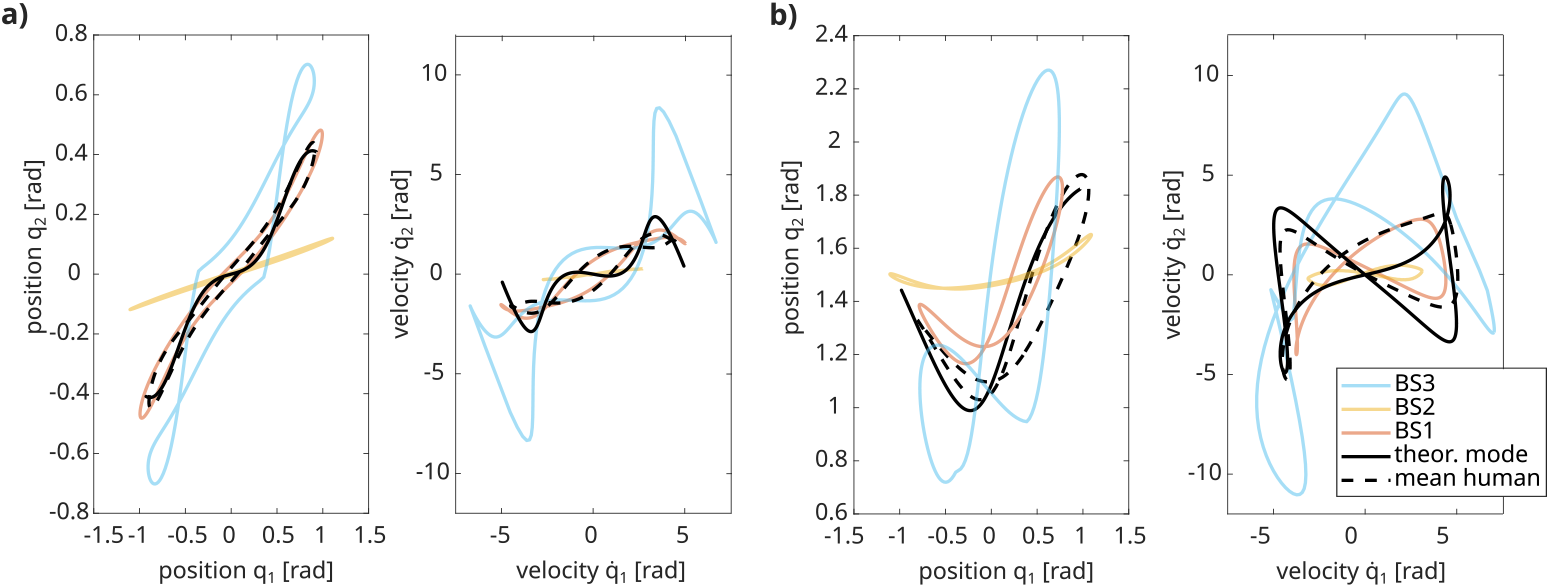
Comparison of the expected baseline strategies with the normal mode for the ideal system (solid black) and the averaged data from the human user study (dashed black). **a)** shows the comparison for the *P0* pendulum configuration in position (left) and velocity (right) space, while **b)** shows the respective plots for the *P90* configuration.

##### Oscillation Frequency

For the first two baseline strategies, *BS1* and *BS2*, the system frequency was given through the control input, i.e., the commanded sine wave frequencies, being *f* = *f*_res_ and 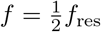 respectively. For the bang-bang control approach (*BS3*) the frequency of *f*_*P*0_ = 0.984 Hz and *f*_*P*90_ = 1.214 Hz was identified for the two pendulum configurations, respectively. Following our hypothesis, it was expected that humans would intuitively excite the pendulum systems close to the respective natural frequencies *f*_res_ only moving the motor link little in comparison to the pendulum links and thus apply a control strategy similar to *BS1*. Indeed, the average frequency with which the double pendulum was excited in the *P0* configuration by the participants was found to be 0.77 Hz, which is very close to the theoretically expected value of 0.78 Hz. Likewise, the identified average frequency that the participants excited in the *P90* pendulum configuration was with 0.92 Hz, thus identical to the theoretical eigenfrequency of the conservative system.

##### Mode Metric

To further investigate the assumption that humans excited the double pendulum motion, a *mode metric η* was introduced, which is based on the Dynamic Time Warping (DTW) principle to calculate the distance between two signals on different time scales [27]. This mode metric was once computed for the position space (*η*_pos_) solely relating the upper joint angle *q*_1_ and the lower joint angle *q*_2_ and additionally applied to the manifold embedded in the state space 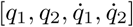 to obtain *η*_man_ since only then the mode is completely defined. With these metrics, the distance between the reference from the conservative case and the resulting pendulum trajectories excited through the baseline strategies or the participants can be computed and compared.

For *BS1* it was found that *η*_pos_(*P*0) = 1.472 and *η*_pos_(*P*90) = 3.153 for the *P0* and *P90* configuration, respectively. Applying *BS2*, the slow control approach, resulted in *η*_pos_(*P*0) = 4.304 for the *P0* configuration and *η*_pos_(*P*90) = 4.885 for the *P90*. The bang-bang strategy as *BS3* led to respective mode metric values of *η*_pos_(*P*0) = 3.024 and *η*_pos_(*P*90) = 6.321. The values of the mode metric *η*_man_ comparing the complete manifold of each control approach showed the same quantitative trends for both pendulum configurations. All values are summarized in Table 1. As expected, for both investigated pendulum configurations, the excitation with the resonance frequency through *BS1* led to the smallest values of both mode metrics *η*_pos_(*P*0, *P*90), meaning that the resulting path in joint space was closest to the computed nonlinear mode of the conservative system. Also, comparing over the complete manifold, *η*_man_(*P*0, *P*90) was lowest for the excitation with the resonance frequency through *BS1*, i.e., motions that were closest to the manifolds of the conservative systems.

**Table 1.**
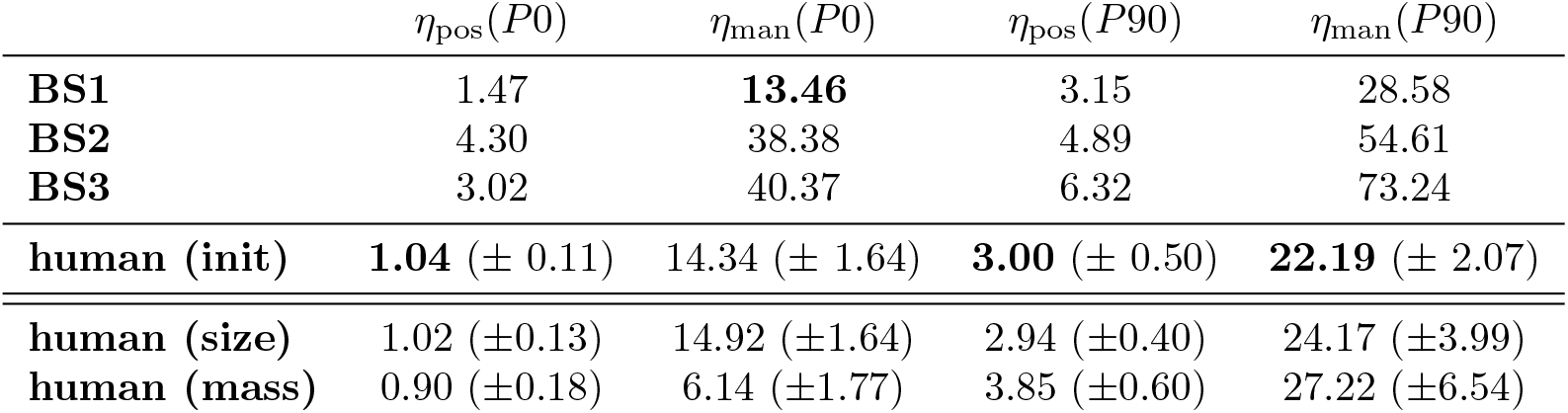
Overview of the calculated mode metrics *η*_pos_ and *η*_man_ for the tested *P0* and *P90* configurations. The first three rows show the results of the identified baseline strategies, followed by the mean value for the human data given with standard deviation over all participants. The respective minimum achieved metric value between these four strategies is marked bold. The last two rows show the computed values for the second experiment run, in which either the target size (*size*) or the lower link mass (*mass*) was altered.

| | $\eta_{\text{pos}}(P0)$ | $\eta_{\text{man}}(P0)$ | $\eta_{\text{pos}}(P90)$ | $\eta_{\text{man}}(P90)$ |
| --- | --- | --- | --- | --- |
| **BS1** | 1.47 | **13.46** | 3.15 | 28.58 |
| **BS2** | 4.30 | 38.38 | 4.89 | 54.61 |
| **BS3** | 3.02 | 40.37 | 6.32 | 73.24 |
| **human (init)** | **1.04** ($\pm 0.11$) | 14.34 ($\pm 1.64$) | **3.00** ($\pm 0.50$) | **22.19** ($\pm 2.07$) |
| **human (size)** | 1.02 ($\pm 0.13$) | 14.92 ($\pm 1.64$) | 2.94 ($\pm 0.40$) | 24.17 ($\pm 3.99$) |
| **human (mass)** | 0.90 ($\pm 0.18$) | 6.14 ($\pm 1.77$) | 3.85 ($\pm 0.60$) | 27.22 ($\pm 6.54$) |

**Table 2.** Parameters for all tested double pendulum configurations investigated in the user study. Only for the variation of Experiment 2, in which the lower link mass was increased, *m*_2_ was changed to 0.625 kg. All other parameters remained.

| upper link |  | lower link |  |  |
| --- | --- | --- | --- | --- |
| $l_1$ | 0.5 | $l_2$ | 0.5 | [m] |
| $m_1$ | 0.5 | $m_2$ | 0.25 (0.625) | [kg] |
| $k_1$ | 5 | $k_2$ | 3 | [N m rad<sup>-1</sup>] |

Applying the mode metric to the human user data, for the *P0* configuration *η*_pos_(*P*0) = 1.04 *±* 0.11 was identified when only considering the position space, and *η*_man_(*P*0) = 14.34 ± 1.64 over the complete manifold. Thus, in position space and in the full manifold space, the values of the mode metric were close to the values obtained with *BS1*. As the nonlinearity of the mode is not so pronounced, the difference between the sinusoidal excitation of *BS1* and the human excitation is not so large. For the flexed *P90* double pendulum configuration, the mode metrics are calculated to be *η*_man_(*P*90) = 3.003 ± 0.50 and *η*_man_(*P*90) = 22.19 ± 2.07 when considering the position space and manifold, respectively. Here, again, the value of the mode metric was very similar to the one obtained with *BS1* in position space. Still, considering the complete manifold, we can clearly observe that the mode metric is considerably lower (by 22%) than for the sinusoidal excitation of *BS1*.

Since the excited frequencies through the participants are for both double pendulum configurations close to the calculated eigenfrequencies, it becomes clear that the participants excited the systems in a similar manner as predicted by *BS1*. The low values of the mode metrics further indicate that the humans indeed exploited the intrinsic dynamics of the double pendulum systems, verifying our stated hypothesis. In the *P0* configuration, where the system’s nonlinearity is not so pronounced, the human’s performance is similar to the one obtained by applying a simple sinusoid with *BS1*. However, for the *P90* configuration, where the nonlinearity of the system is very pronounced, humans clearly outperform the excitation with a simple sinusoid of *BS1*, which is also visually apparent (Fig. 4).

#### Applied handle motion

Exploiting the intrinsic dynamics of the double pendulum for the given task means mainly holding the input handle static and only moving minimally around an equilibrium position. This becomes apparent when plotting the handle coordinate *θ* controlled by the human user through the haptic input device against the upper link coordinate *q*_1_ for the *P0* (Fig. 5a, right) and *P90* (Fig. 5b, right) configuration. For both cases, it can be seen that the input handle was moved little compared to the amplitude of the pendulum link. This fits the prediction through *BS1*, as the system mainly moved through the energy stored in the springs, while the human input was primarily necessary to compensate for friction and control the energy level. Comparing the overall motion applied to the motor link averaged over all participants results in a sine wave-like motion that seems to have a phase lag of around 0.5*π*, which coincides with the results of *BS1* exciting the systems with their respective eigenfrequency.

**Fig 5.**
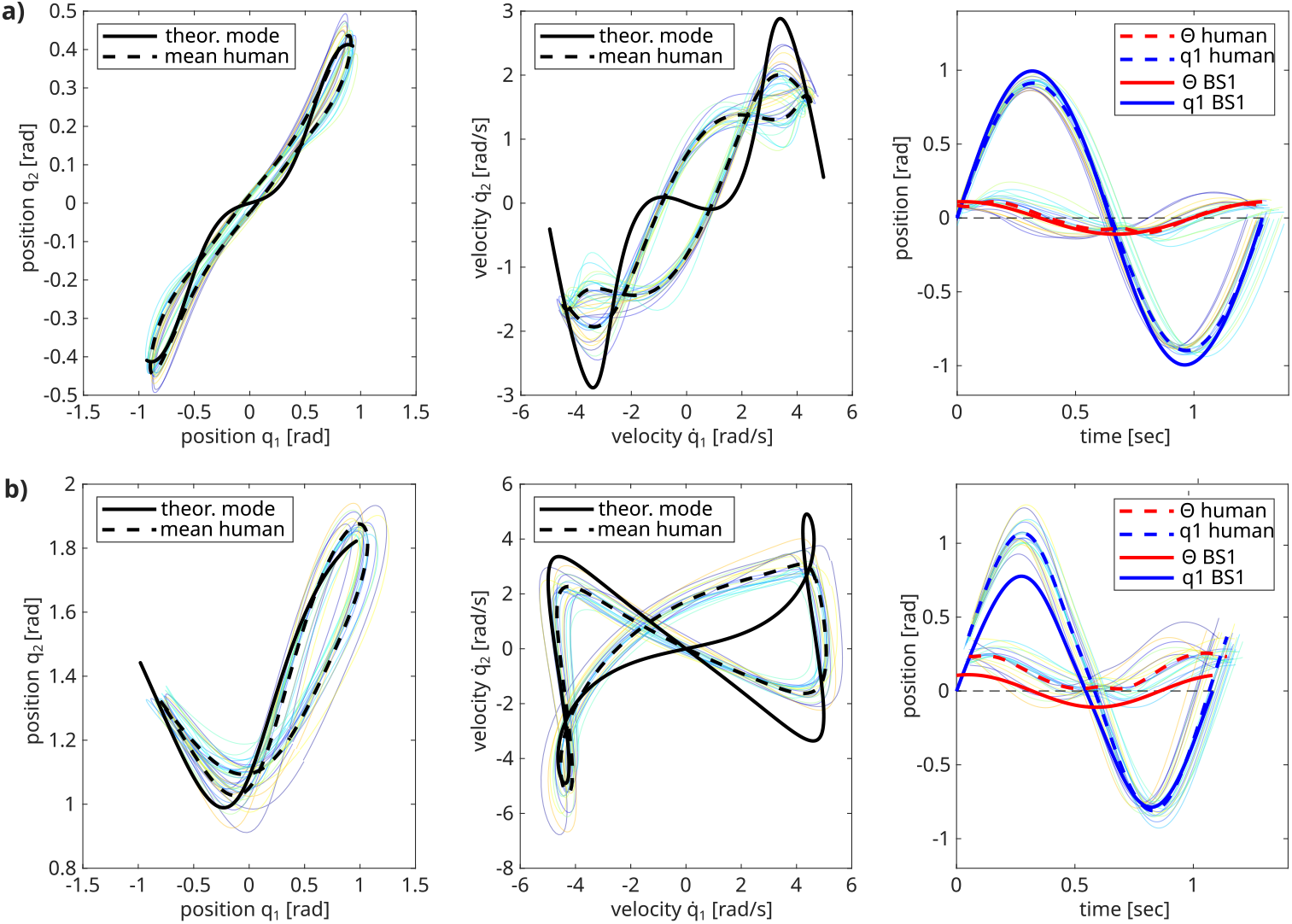
Comparison of the mean pendulum motion per period of all individual participants (colored) overlayed with the averaged resulting trajectory of all participants (dashed black) for **a)** the *P0* pendulum configuration and the **b)** the *P90* configuration. The left plots show the trajectory in position space, the middle plots in velocity space. The black solid lines show the computed modes for the ideal systems. On the right plots, the motion of the red handle that the participants commanded is related to the motion of the upper pendulum link coordinate *q*_1_ (blue). Here, the dashed lines refer to the participant data again, while the solid lines relate to *BS1*, in which the system was excited with its resonance frequency.

However, the *P90* configuration shows an apparent offset to the handle position that participants applied compared with the expected linear theory. Instead of oscillating the pendulum around the zero-position as defined in Figure 2a, the participants shifted the handle to an average value of *q*_1_ = −7.44° = −0.13 rad thus shifting the equilibrium position of the spring for the upper pendulum link. This behavior was constant throughout all participants.

Taking a closer look at the individual curves per participant (Fig. 5a and b, right), however, it is visible that the particular strategy on how exactly the handle was moved varied clearly for the different participants. The first thing to notice is that most participants’ input plateaued at the zero-crossing, meaning they stiffened in their position to hold the input link. Overall, two different main strategies can be identified, where the participants either are in phase with the pendulum, thus pushing it after the zero-crossing, or in anti-phase to take out energy while the pendulum is still moving toward its target. In fact, the observed phase lag of the handle motion compared to the upper link motion varied between 0.15*π* and 0.70*π* for the stretched *P0* configuration. Similarly, for the flexed *P90* configuration, a variation in phase lag of 0.21 − 0.74*π* was found. Nevertheless, although the applied phase shift of the motor link varied widely, the resulting amplitude difference between the motor and pendulum links remained very consistent throughout all participants (Fig. 6a). Comparing the applied phase lags of each participant individually for the two pendulum configurations shows that most participants were consistent in their chosen strategy independent of which system they were excited. This indicates that although all participants found a way to exploit the intrinsic dynamics of the two double pendulum systems (Fig. 5 left and middle), the individual strategies to move the handle varied clearly.

**Fig 6.**
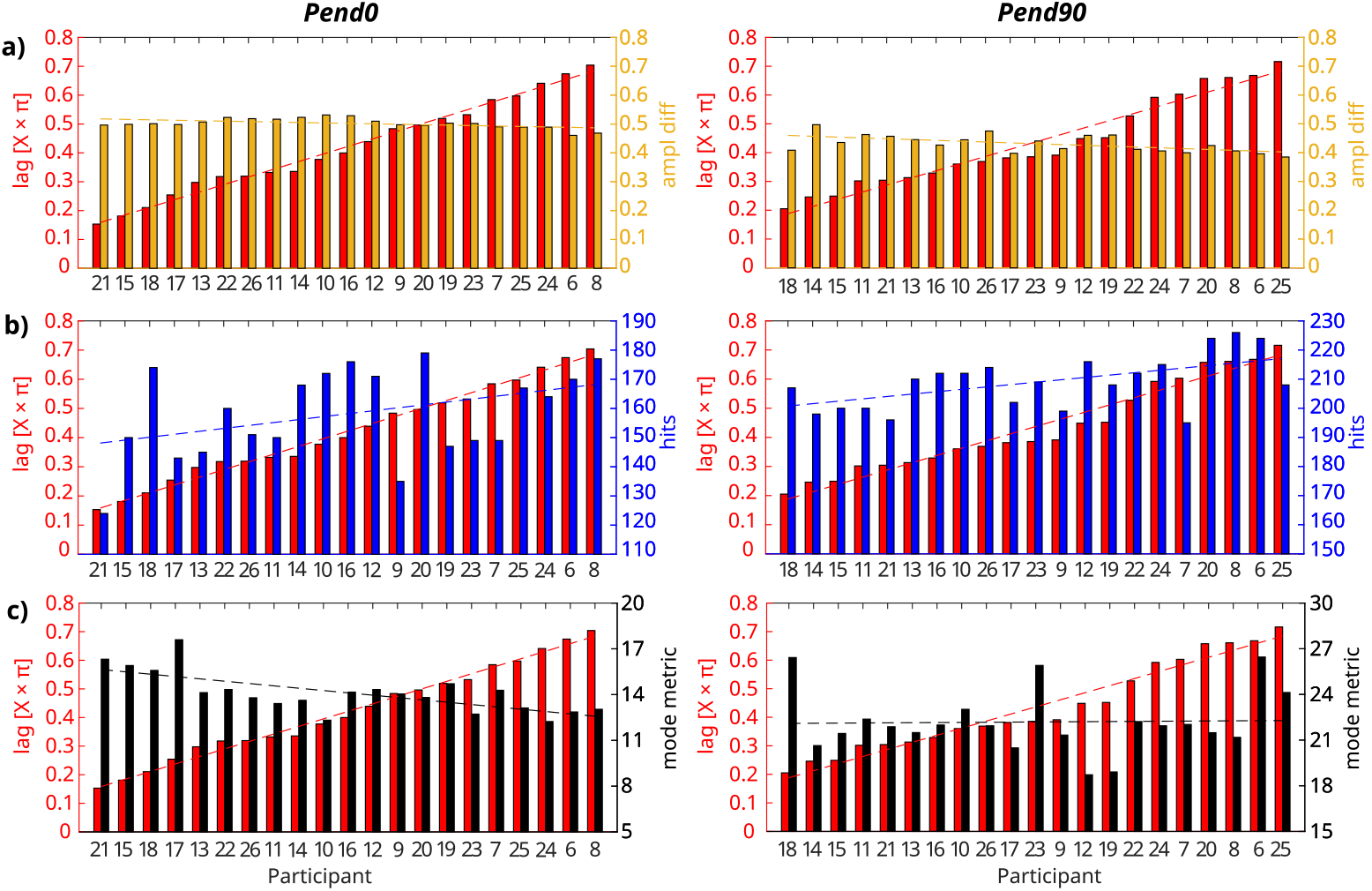
Trend comparison of identified metrics per participant during the human user study with the *P0* configuration on the left and the *P90* configuration on the right. The x-axis indicates the participant number. Shown is the phase lag of the motor link commanded by the participants relative to the upper pendulum link (red) in comparison with the **a)** resulting mean amplitude difference between the pendulum link and motor link, **b)** obtained hits per participant over the three best trials and **c)** computed mode metric over the complete manifold. Dashed lines indicate the linear fit through the bar data.

To further investigate whether the individual strategy chosen by the participants affected the performance, the applied phase lag per participant was compared to the obtained hit score (Fig. 6b). Sorting the individual performance by hit score and comparing it with the respective applied phase lag applied by each participant shows for both configurations a trend line suggesting a relationship between the applied phase lag and the obtained hit score. Computing the correlation between the two variables for each configuration shows indeed a significant correlation for the *P90* configuration (*p* = 0.01), while the significance level is not quite reached for the *P0* configuration (*p* = 0.09). Especially in the *P90* configuration, the three participants with the highest scores (Participant 6, 20, and 8) applied a similar phase lag of 0.6*π*, which was also seen in the *P0* configuration for Participant 6 and 8. This could suggest that a phase lag in this range could be advantageous. However, it is contradicted by the data of, e.g., participant 20, who applied a much lower phase lag strategy in the *P0* configuration while still achieving a high hit score. Carrying out a similar comparison between the phase lag and the mode metric (Fig. 6c) shows a clearly significant correlation for the *P0* configuration (*p* = 0.001), indicating that the highest phase lag of around 0.6*π* led to the lowest values for the mode metric. This would mean that the excited system was closest to the expected idealized eigendynamics, suggesting that the system’s compliance was best used. However, this trend cannot be found for the *P90* configuration. In fact, Participant 6, who was under the top 3 highest-scoring users, had one of the highest values of the mode metric in the *P90* configuration. From the obtained results, it seems that lag alone is not a clear predicting factor neither for the successful execution of the task nor for how well participants exploited the eigendynamics of the systems, and again individual skill and preferences also play a role.

#### Task success

Although the actual score reached by the participants was only of secondary interest, it was still considered as a metric for how successful the participants achieved the given task in the high-score game. For the *P0* pendulum configuration, the participants averaged 158 ± 15 hits over their three best trials. In an ideal trial where no overswing or error occurred, the maximum possible score would have been 187 with the identified eigenfrequency of each system. This means that the participants hit the target with an accuracy of 85%. With the flexed pendulum in the *P90* configuration, the participants averaged 209 ± 9 hits over their three best trials compared to an ideally possible score of 221 hits when excited with the ideal eigenfrequency. Thus, the participants’ hit performance was actually better with hitting the target in 94% of the approaches.

### Experiment 2

The results of the second experiment, in which either the ball size was decreased or the lower link mass was increased, emphasized the consistency of the participants. First, investigating the scenario with the decreased target size, it was expected that the participants might alter their control strategy to move much slower since a higher accuracy was required to hit the targets without overswinging. Indeed, the excited system frequency was significantly lower when the target size was decreased, leading to average values of 0.75 Hz and 0.87 Hz for the *P0* (*p* = 0.01) and *P90* (*p* = 0.005) configuration, respectively. The decreased frequency due to the higher required accuracy resulted in a significantly lower number of achieved hits in both configurations with a hit rate of 0.74% and 0.87% for *P0* and *P90*, receptively. Thus, for each pendulum configuration, the hit rate dropped slightly. Although showing statistical significance, the values of the excited frequencies are only marginally lower than in the initial setup, and the still low values of the mode metric (Tab. 1) indicate that the excited pendulum motion was similarly close to the ideal reference as with the initial target size (Fig. 7a and b). Thus, despite the slight drop in oscillation frequency, overall, the participants still exploited the intrinsic dynamics of the pendulum systems and generally applied the same strategy (Fig. 7, right plots), proving to be very robust in their excitation approach.

**Fig 7.**
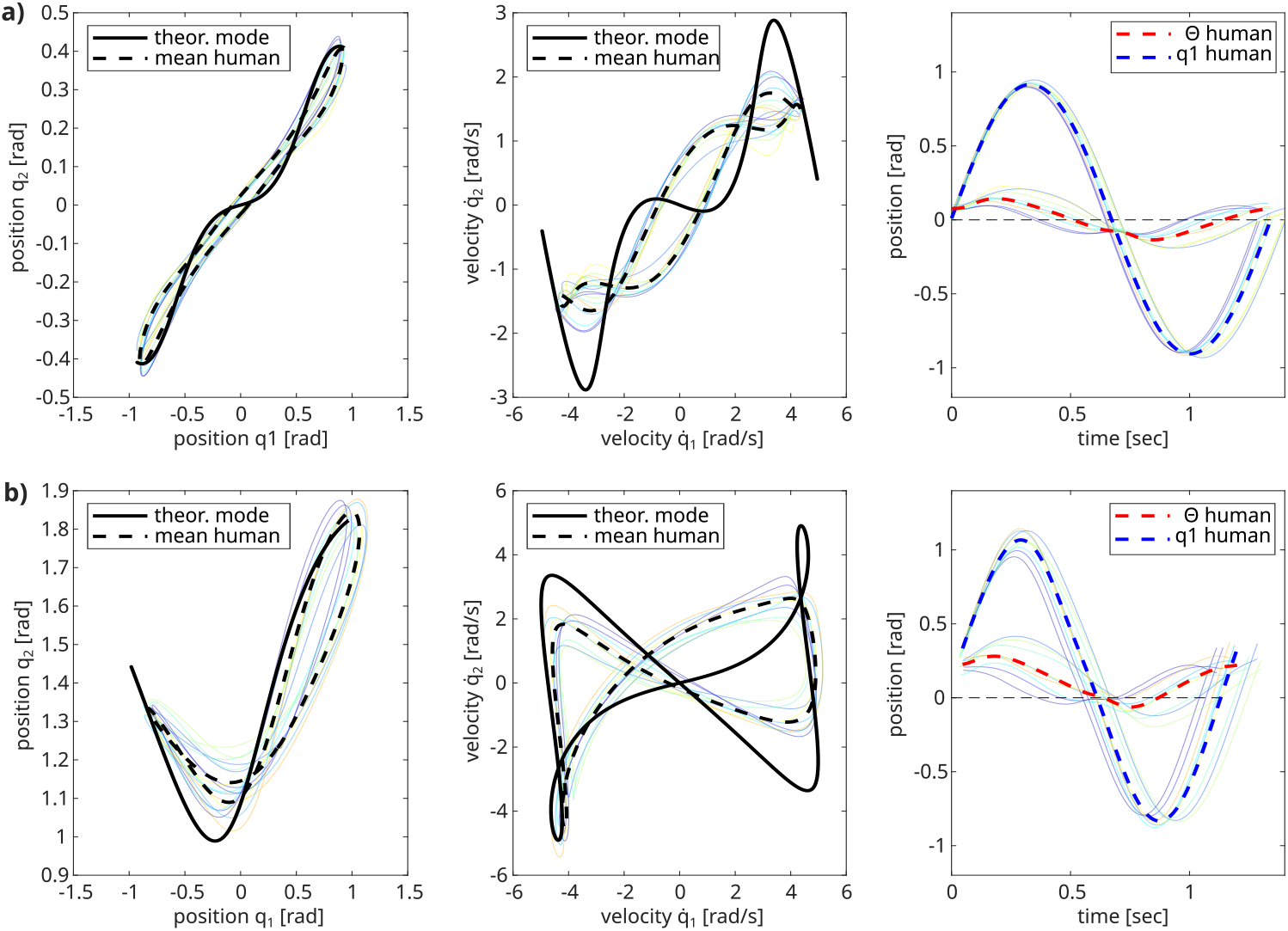
Comparison of the averaged participant data (dashed) with the computed modes (solid black) for the **a)** *P0* and **b)** *P90* configuration obtained in the experiment run with decreased target size. Colored plots show individual participant data. The right plots compare the red handle coordinate *θ*, and the upper link coordinate *q*_1_ (blue).

In the second group of the experiment variation, the mass of the lower pendulum link was increased, but the target size stayed identical. Since the system was altered, the corresponding nonlinear modes were also expected to change. However, it was expected that the participants would still apply the same strategy and exploit the intrinsic dynamics of the system. The calculation of the altered modes showed that the ideal systems would oscillate with a frequency of 0.52 Hz and 0.64 Hz in the *P0* and *P90* configuration, respectively. As expected, the participants again hit close to this resonance frequency with 0.53 Hz and 0.65 Hz. Comparing the achieved hits with the ideally possible ones at this frequency, the participants could actually achieve slightly better hit rates as in the initial setup, resulting in 0.88% and 0.98% for the *P0* and *P90* configuration, respectively. This is likely due to the slower-moving system giving the users more time to evaluate the anticipated motion. Our hypothesis is once more validated by the mode metrics (Tab. 1) that actually showed that the excited motion was closer to the theoretically expected one for the *P0* configuration and in a similar range for the *P90* configuration as also visually apparent in Figure 8a and b, respectively. The detailed comparison of the handle motion coordinate *θ*, and the upper link *q*_1_ also showed a very similar pattern to what was observed for the initial setup.

**Fig 8.**
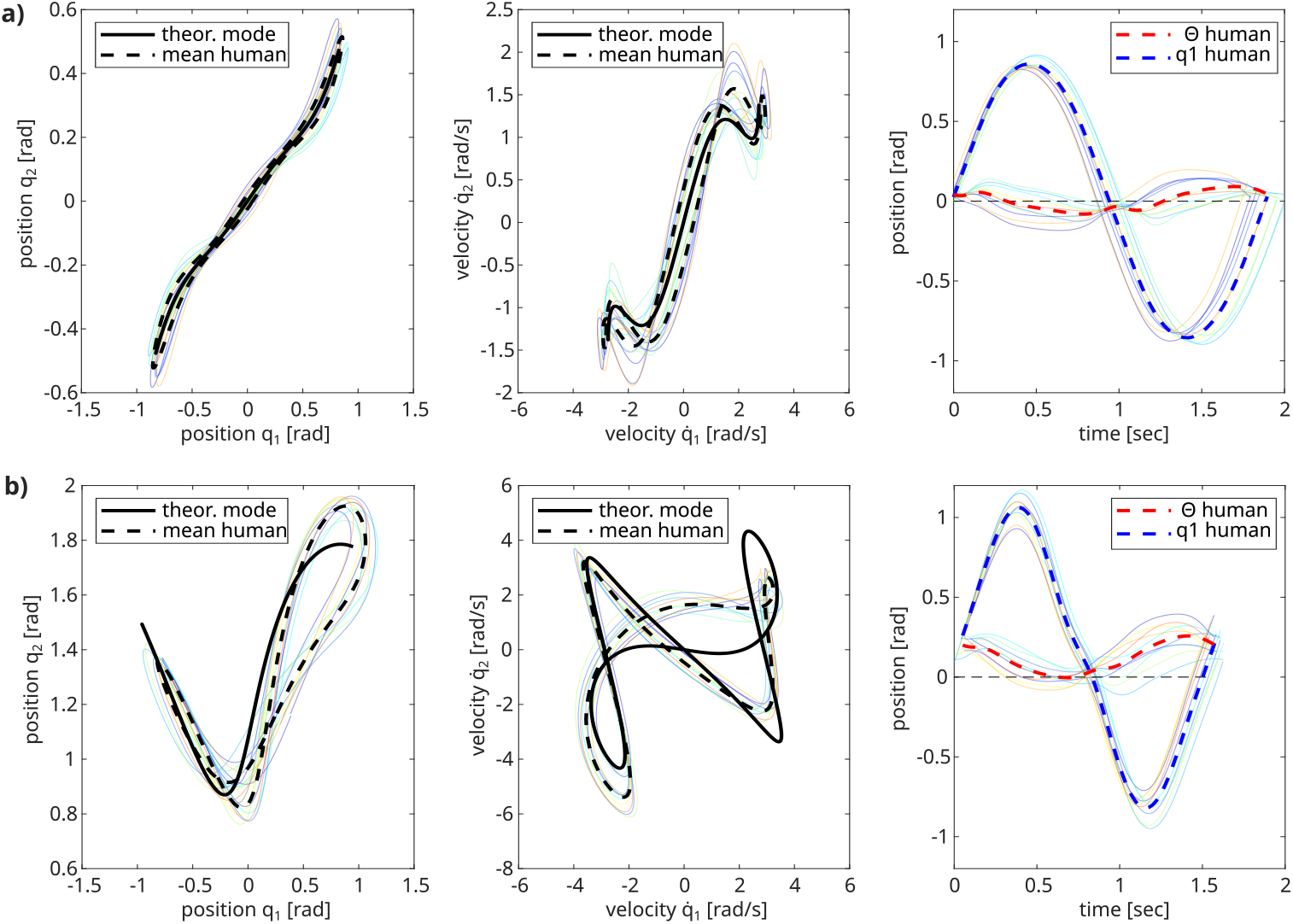
Comparison of the averaged participant data (dashed) with the computed modes (solid black) for the **a)** *P0* and **b)** *P90* configuration obtained in the experiment run with increased lower link mass. Colored plots show individual participant data. The right plots compare the red handle coordinate *θ*, and the upper link coordinate *q*_1_ (blue).

These findings further support our hypothesis that humans intuitively adapt their control strategy to exploit intrinsic motions and can also very successfully scope the dynamics of different mechanical system.

### Experiment 3

The previous experiment results showed clearly that humans can estimate the behavior of systems with complex dynamics and use this knowledge to exploit the dynamics in periodic tasks. However, in the previous experiment setup, the given task was to oscillate between two points that were located along the mode of the system, i.e., providing ideal conditions to make use of the system’s intrinsic dynamics. From these findings, the question arises as to how a human will excite a system in the same task when the targets are not directly located on the eigenmode, such that only supporting the system’s intrinsic motions will not enable the user to hit the targets. Thus, an additional experiment variation was conducted, where a double pendulum configuration was tested with target balls that were not located along that system’s mode. In this variation, the spring offset between the upper and the lower link was set to 45 ° (*P45*), thus representing an intermediate configuration between the already described *P0* and *P90* configurations. However, for this case, the balls were not located within the endpoints of the nonlinear mode computed with the mode tool (Fig. 9a), but instead arranged to lie on a radius between the targets for the two previously described system configurations *P0* and *P90* (Fig. 9b and c).

**Fig 9.**
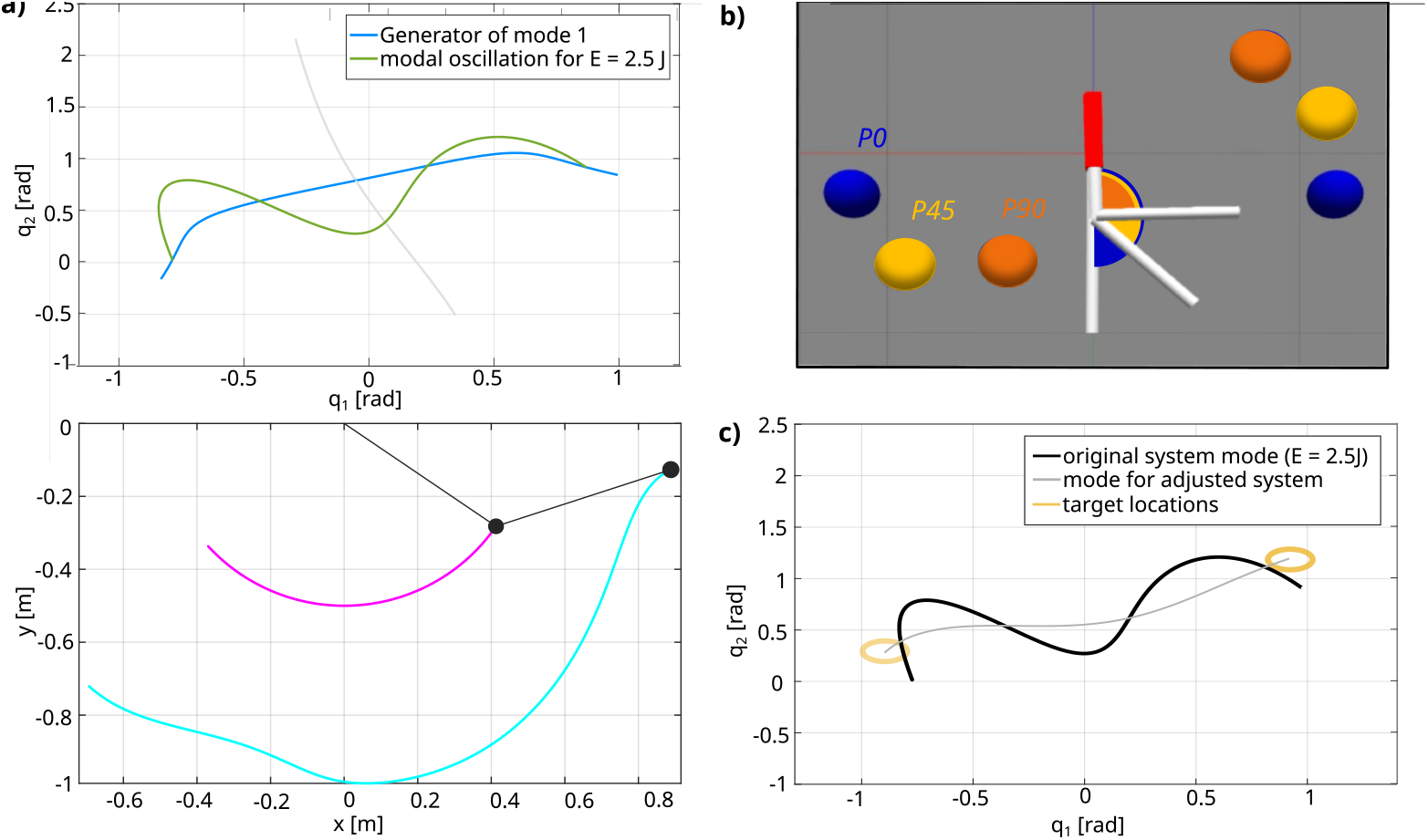
**a)** Computed mode for the *P45* pendulum configuration with the initial parameters from Tab. 2 in position joint space (top) and Cartesian space (bottom). **b)** Target location for the *P45* configuration (yellow) arranged between the targets of the *P0* (blue) and *P90* configuration (orange). **c)** Comparison of the computed mode for the *P45* system (black) and a reverse-engineered mode with altered parameters in this configuration (grey).

For further analysis of the results, the *mode tool* was additionally used to reverse-engineer a double pendulum system in the *P45* configuration for which the chosen target locations would indeed be the turning points of its mode. It was computed that by altering the lower link mass to *m*_2_ = 0.125 kg and the spring between the two links to *k*_2_ = 2.5 N m rad^−1^, the turning points of that system’s mode would indeed lie withing the chosen target locations for an energy level of 2.5 J. The eigenfrequency for this altered configuration is 1.04 Hz. The comparison of that newly computed mode with the one for the original *P45* parameters that were used in the experiment can be seen in Figure 9c). The mode of this altered double pendulum system is taken as an additional reference to compare the obtained results of the participants.

Although the target balls were not located on the modes, the participants were intuitively able to excite the pendulum system to alternately hit the target balls. In fact, there was no drastic drop in performance noticeable, judged by the achieved hit points, where they reached an average of 143 ± 9 over their three best trials compared to a maximum of 197 ideally achievable. Thus, the participants had a 72% hit rate, which is in the same range as seen for the previously investigated pendulum configurations. Again, the identified frequency of the pendulum was with 0.81 Hz close to the computed eigenfrequency of 0.82 Hz for the ideal conservative system analyzed with the *mode tool*. However, since the targets were not located within the endpoints of the modes, the resulting pendulum motion cannot and does not correspond to the expected theoretical mode (Fig 10c). The mode metric calculation led to values of *η*_*pos*_ = 4.22 ± 0.16 in the position space and *η*_*man*_ = 71.91 ± 1.34 considering the whole manifold. Specifically, the latter value indicates that the excited pendulum motion was quite different and thus “far” from the ideal intrinsic motion as also apparent in Figure 10. Nevertheless, it is very noticeable that throughout all participants, the excited pendulum motion was very consistent, and the mean variation between the participants was in the same magnitude range as for the *P0* and *P90* configuration.

**Fig 10.**
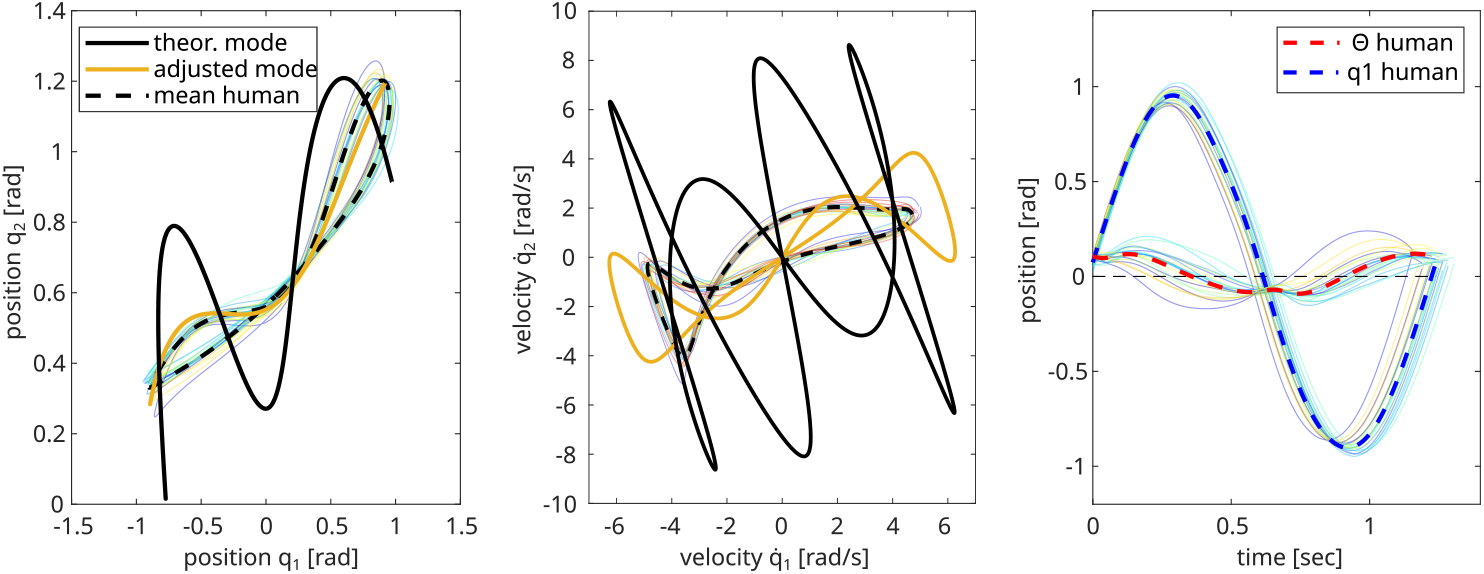
Comparison of the excited period pendulum motion averaged over all participants (dashed black) for the *P45* pendulum. The left and the middle plots show the joint positions and velocities of the trajectory, respectively. The black solid line visualizes the mode of the ideal system with the initial parameters from Table 2, while the yellow mode trajectory was computed for a system with altered mass and stiffness. The right plot compares the applied participant handle motion (red) with the excited upper pendulum link trajectory *q*_1_ (blue).

Examining the handle motion closer, it becomes apparent that the participants still chose the same overall excitation strategy for the pendulum system as seen for the two previously presented systems, i.e., mainly holding the handle still and only minimally moving to compensate for energy losses and trigger direction changes. Thus, as before, the amplitude of the pendulum motion is comparably big to the handle motion, thus still making use of the system’s compliance (Fig 10, right).

Comparing the excited pendulum trajectory of the participants with the reverse-engineered system where the targets would indeed be located at the endpoints of the mode shows a surprising similarity (Fig 10, yellow lines). Computing again the mode metrics of the participant data against this mode, shows indeed considerably lower values for *η*_*pos*_ = 1.90 ± 0.16 and *η*_*man*_ = 27.06 ± 1.06, which are in fact similar to the mode metric values that were obtained for the *P0* and *P90* configurations (Tab. 1).

## Discussion

A human user study was carried out to test the hypothesis that humans can scope and exploit intrinsic motions of mechanical systems, even when those systems exhibit clearly nonlinear dynamics. In this study, participants were asked to excite different configurations of a virtual compliant double pendulum through a haptic joystick. The joystick’s position commanded a motor input link that was coupled to the upper pendulum link through a spring. The resulting forces of that spring were reflected to the participants, i.e., giving the impression of holding and exciting a flexible stick. The task was to excite the system to alternately hit two targets with the goal to precisely hit the target as often as possible in a set time frame of 40 s. Enabled through a newly developed *mode tool* that can compute nonlinear normal modes, i.e., expected intrinsic modal oscillations a system is mechanically inclined to do in the absence of friction, we could predict how the excited pendulum system should move when optimally exploiting its eigendynamics.

### Humans exploit elasticities and intrinsic motions

In the first experiment part, the targets to hit were located on the turning points of these predicted motions, and the excited pendulum motions were compared to the idealized expectation. In this setup, two double pendulum configurations were tested with a spring offset between the upper and lower link, either being 0 ° (*P0*) or 90 ° (*P90*). Minor friction was added to the system to give the participants a control objective. After some minutes of initial training to explore the system and get used to the setup, all participants intuitively chose an excitation strategy for both pendulum configurations that excited the system in its intrinsic dynamics, i.e., along its respective eigenmode. This was verified through the comparison of multiple metrics:

First, the oscillation frequency of the pendulum systems excited through the participants matched very well the theoretically computed values of the expected eigenfrequencies in the conservative case.

Second, the DTW-based *mode metric* was used to quantify the match of the excited pendulum motion through the participants to the ideally expected mode motion. Through this metric, the excited pendulum motions and the ideal reference were compared once solely considering the position space and, additionally, taking into account the complete manifold, which fully defines a mode. To set the resulting values of this metric into perspective, the obtained values from the participant data were compared to the three baseline strategies that were defined prior to the experiment, i.e., a sine wave motion with either the systems’ eigenfrequency (*BS1*) or a much slower frequency (*BS2*) and a bang-bang controller (*BS3*) derived from previous robotic experiments [23]. As expected, the mode metric was lowest when the systems were excited at their respective eigenfrequency, which was the case for *BS1* and the strategy applied by the participants. This means that the resulting path and manifold of the double pendulum systems were closest to the theoretically predicted ones, which exploit the eigendynamics. Comparing these two strategies, the mode metric values were close for the *P0* configuration, while for the *P90* configuration, the participants outperformed the sine wave excitation.

Next, the difference of the mean amplitude per period of the motor input link commanded through the participant (or controller) and the excited upper link motion was compared to further quantify if the elasticities of the systems were exploited. Indeed, for *BS1* exciting the systems with the eigenfrequencies, the upper link amplitude was 0.56 ° larger than the commanded motor link for the *P0* configuration and 0.41 ° for *P90*. Similarly, the participants’ excitation strategy led to a mean amplitude difference of 0.49 ± 0.03 rad and 0.43 ± 0.04 rad) for *P0* and *P90*, respectively.

Summarizing, we present first data-based evidence that humans can indeed intuitively scope and support systems in their nonlinear dynamics, verifying our initial hypothesis. This is shown with different double pendulum systems, which were intuitively excited at their eigenfrequency, and the pendulum link motions were much larger than the commanded motor link positions, i.e., the systems’ elasticities were exploited. These findings were consistent throughout all participants. Additionally, the defined mode metric shows that the resulting pendulum motion excited through participants is close to the theoretically expected paths and manifold. The findings also indicate that the higher the system’s nonlinearity, the better the human gets compared to a sinusoidal excitation baseline.

Multiple reasons can be identified as to why humans choose to exploit the inherent dynamics of the mechanical system in the given setup:

1. Since the task was to hit a target as often as possible while giving some leeway on the accuracy, the human could make the expected trade-off according to Fitts Law [28] between accuracy and speed in favor of the latter. If accuracy had been a stronger requirement in the experimental task, participants would have been expected to move more slowly, e.g., as shown in *BS2*, where the slow sine wave drove the motor link. However, since the focus was on maximizing points, requiring faster movements, this strategy was only applied by some subjects in the beginning to explore the system dynamics.
2. When focusing on speed, the humans could have chosen a method where they move fast with the system. However, this would mean the speed of the pendulum motion needs to be precisely matched to avoid adding energy to the system, which in turn would destabilize the system motions. Learning such a velocity profile is hard and, in the given scenario, also reaching the limits of human capabilities. In threat situations, humans can perform reaching tasks with around 200 − 300 ms and maximum velocities of up to 2.5 m s^−1^ [29]. In the slowest case of the experiment setup (*P0*), the eigenfrequency was 0.78 Hz, meaning one back-and-forth motion between two targets should not take longer than 1282 ms including two turning points. Additionally, it needs to be noted that the above-stated values were obtained in a threat situation and that the human does not reach such high velocities in shorter distance motions [30]. Even if continuously moving at this speed in a periodic oscillation would be physically possible, it would require a very high mental workload.
3. Research has shown that humans apply energy-minimizing strategies in interactions with objects [31, 32]. Generally, it is believed that energy-minimization might be one driving factor during human (loco)motions, often being used as one of the cost functions in human-inspired control strategies [33–36]. Consequently, there are multiple reasons that explain why humans generally might strive to exploit the intrinsic dynamics of compliant systems whenever possible in line with the task requirements.

Nevertheless, applying a strategy that supports a system in its intrinsic dynamics leaves the prerequisite that the human must also be capable of exploring and matching the system dynamics. This skill is mainly learned through experience with different objects and extrapolated for different interaction scenarios [5, 12, 13, 37]. In particular, visual cues are known to play a key role in estimating system dynamics. As humans favor strategies that make interactions predictable [2, 7, 8], it seems to be the obvious choice to exploit the intrinsic dynamics if the human can explore the dynamics of a system sufficiently well. Recent research by Razavian et al. [38] has shown that even without visual feedback, humans are able to learn the system dynamics of complex objects, extending previous work showing that novel dynamics can be learned in the absence of visual feedback [39, 40]. Thus, it would be intriguing to further investigate the human capabilities to excite nonlinear dynamics with the presented methods in further experiments where visual feedback is occluded to better quantify these abilities.

### Humans are robust in their control strategy

In further experiments, either the target size was decreased to require more accuracy from the users, or the mass of the lower pendulum link was increased to alter the system dynamics. The results of these experiments showed that humans are very robust in their control approach. Neither of the experiment alterations led the participants to noticeably modify their control strategy. After just a short initial re-training, the participants applied the same identified baseline strategy, showing the same overall features as for the initial setup. However, for both alterations, it was observed that the excited frequency differed slightly from the ideally expected ones (0.01-0.05 Hz), which suggests that the participants were less stiff in their arm position, such that the combined frequency of the overall system could be slightly altered. Although the hit rate dropped about 10% when decreasing the target size, a relatively high hit rate of above 70% could still be maintained. Similar hit rates as in the original setup could be observed with the initial target size but altered masses and, thus, dynamics. These additional experiments indicate that the participants did not, by chance, resort to the applied excitation strategy but strategically exploited the system elasticities and can do so for different systems. The fact that the strategy was not altered for the smaller target size suggests that the chosen strategy still seemed most promising to the humans for achieving the given task, accepting the lower hit rate as a trade-off for speed, in line with Fitt’s Law [28]. Accordingly, the participants might likely alter their strategy when even higher precision is required. This is also suggested by the work of Nagengast, Braun, and Wolpert (2009), which showed that human control strategy in interactions with objects with internal DOFs can be accounted for by a simple cost function representing a trade-off between effort and accuracy [14]. Following these observations, our findings suggest that for a task with higher precision, a more careful excitation strategy similar to *BS2* would be chosen, although this requires further experiments to investigate the relationship between the selected excitation strategy and the specific task goal.

### Humans adapt to individual skill level

Overall, our results show that the human users exploited the elasticities of all the tested double pendulum systems. However, taking a closer look at the individual strategies in comparison with the identified baseline *BS1* gave additional insights. Although all participants applied a strategy in which the motor handle link had to be moved noticeably less than the excited upper pendulum link by exploiting the system elasticities, the phase lag between the two links varied widely per participant (Fig. 6a) This indicates that while it seemed intuitive for all participants to exploit the eigendynamics, how exactly this was achieved (i.e., at which moment energy was added to the system) depended on the participants’ individual preferences and capabilities.

The experimental data suggests that a phase lag around 0.6*π* might be the best choice to most efficiently excite the system and obtain a high score. However, although these observations are intriguing, the present data is inconclusive, and further experiments should be carried out in which the friction is varied to investigate whether participants converge to a common strategy with a similar phase lag value.

When explicitly comparing the strategy applied by the participants and the comparable reference approach of *BS1* in the *P90* configuration, another important note is made: Although the achieved overall amplitude difference for this configuration was almost identical (*∼*0.4 °) for the participant strategy and the baseline (Fig. 5b), in the participant data, a clear shift of the chosen zero position for the handle could be observed. While in the theoretical case and also for the baseline strategy, the zero position of the motor link was set perpendicular to the mounting point as shown in

Figure 2a, all participants shifted the point around which they oscillated to *θ*_0_ = −7.44° = −0.13 rad, thus moving slightly towards the right ball (Fig. 13b, orange). One possible reason for this might be a distorted perceived symmetry. Although the endpoint of the double pendulum in the *P90* configuration is closer to the right point than the left, the handle position lets it appear as if the system is closer to the left target. Thus, the participants might have shifted the handle slightly to the right to compensate for this feeling. Since the target size gave some leeway to the accuracy, this shift might have actually slightly altered the mode, leading, in this case, to an improved performance judged by the mode metric compared to the respective baseline strategy. Once more, this observation needs to be investigated in further experiments to provide conclusive data.

### Humans exploit intrinsic motions outside modes

Finally, experiments where the targets were not located on the mode show that, in this case, the excited pendulum motion does not coincide with the intrinsic system mode for the same energy level as for the *P0* and *P90* configuration. This is expected since the targets were not located on the modes so it would not make sense to excite the system along the idealized mode. Instead, the humans adapted their strategy to be able to hit the targets as required by the task. Nevertheless, comparing the handle motion to the upper pendulum link motion (Fig. 10), it becomes obvious that the elasticities of the system are still being exploited since, again, the handle is only moved little compared to the pendulum. This indicates that humans still try to exploit elasticities in compliant systems when the task motion is not in line with the intrinsically preferred motion of the system. This is reasonable since, in everyday life, tasks might not always be possible through the excitation of a system’s mode motion. However, it is likely that even when not the ideal motion of the system is followed, humans still make use of the inherent dynamics, which is supported by the fact that the excited oscillation frequency was, even in this experiment, very close to the ideally computed one. Thus, the dynamics were still used but altered, e.g., by inserting energy at different moments and/or lowering the applied energy. Therefore, the experiment supports the findings that humans have very advanced capabilities to estimate the intrinsic dynamics of compliant systems and will exploit the elasticities when possible. Doing so, they seem to be able to shape the system motion, probably by altering the stiffness of their limb accordingly [41–43]. A possible explanation of the behavior comes from the area of robot motion control, from the concept of controlled Lagrangian [44, 45]. There, the robot controller slightly changes the closed-loop dynamics of the system to match desired tasks, while the resulting dynamics still correspond to the mechanical system. Further experiments, including EMG measurements, would be necessary to explicitly address this hypothesis.

## Conclusion

The presented research could, for the first time, provide data-based evidence that humans can estimate intrinsic motions of compliant systems, even with nonlinear dynamics, and intuitively choose a control strategy that matches and exploits these. Thus, the contribution of this research is twofold: first, we presented a novel method to better investigate how humans might choose their excitation strategies in everyday interactions and, second, the findings provide first insights about the human capabilities to estimate and adapt to the dynamics of external systems. Although further research is needed to investigate the chosen human strategies in more detail and in more complex scenarios, this research could help to reveal some of the underlying mechanisms driving human motion planning and control during the excitation of periodic motions, which in turn might lead to generation of new hypotheses on a neural level.

## Methods

### System implementation and visualization

The double pendulum systems the participants interacted with during the human user study, were implemented in a virtual environment using Gazebo 11. The set parameter values of the systems were loosely based on the dimensions of a human arm and are detailed in Table 2. The values are identical for all measurements, except a variation in Experiment 2, where the mass of the lower pendulum link was increased to 0.625 kg. The default ODE solver was used to integrate the system dynamics. The system was driven through a motor link, which was implemented as a visual additional link with the coordinate *θ* (Fig. 11, red link). This link was connected to the upper pendulum link with a spring of stiffness *k*_1_, such that the position of the motor link defined the equilibrium position of the double pendulum system. In simulation, the motor link could be directly commanded by position control. This torque was reflected as force feedback to the 1-DOF joystick (Fig. 11). The corresponding values were real time computed and commanded in C++. By utilizing the received force-feedback the user was given the impression to hold a real object in their hand and shaking it to excite sustained oscillations like in a natural interaction. Since in real life scenarios energy losses always occur due to internal and external effects, friction was added to both pendulum joints with the values of *d*_*i*_ = 0.02*k*_*i*_. This made a sustained control action of the human users necessary to sustain a motion of the system with a constant energy like that showcased by the example of a swing on the playground. Thus, the users had to find a strategy to sustain the pendulum motion to fulfill the task of alternately hitting two target positions in the experiment. The setup and task definition will be described in more detail in the following section.

**Fig 11.**
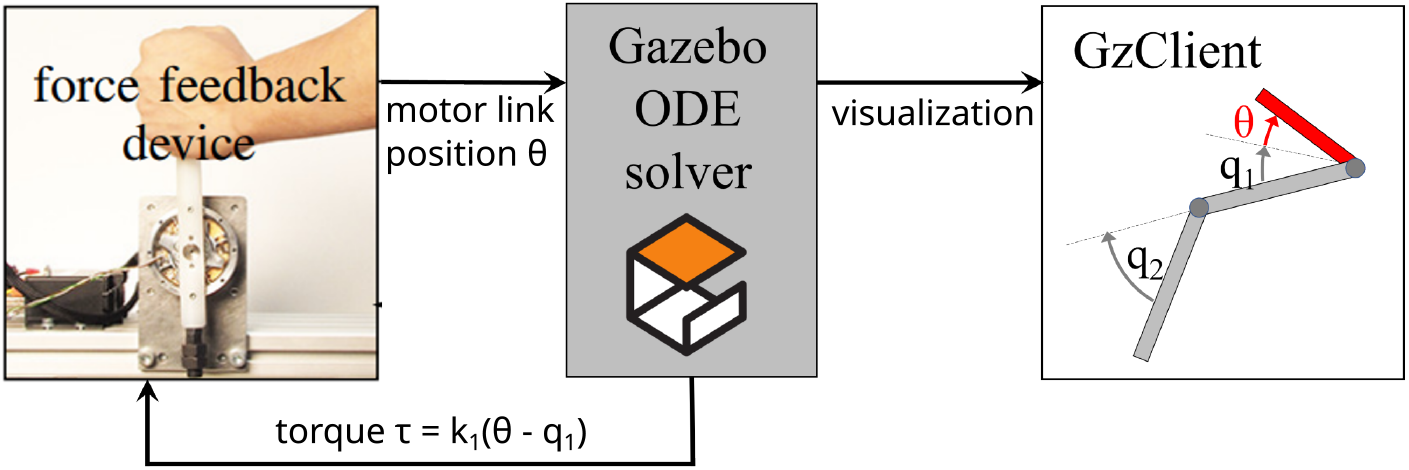
The human participants command the virtual double pendulum systems implemented in Gazebo through a haptic joystick interface that maps to the position of a motor link (*θ*, red), which is connected to the upper pendulum link (*q*_1_) by a spring. The spring forces *τ* are reflected to the user as haptic feedback through the joystick.

The targets to hit with the pendulum were implemented as links with ball geometry in Gazebo. To visually indicate to the participants that they had sufficiently reached the target to earn a hit point, a custom Gazebo plugin was implemented that changed the color of the target balls whenever a collision with the lower pendulum link was detected.

The different software implementations were interfaced through the DLR-intern *Links and Nodes*. The control loop was running at 1 kHz. The angle velocities of all joints were recorded as well as the forces reflected to the user and the information displayed to the user on screen.

### Characterization of system behavior

Prior to the human user study, the two investigated configurations of the double pendulum system *P0* and *P90* were characterized in simulation to render the behavior of the systems without a dedicated control strategy and outline the reachable space. For this purpose, first a sweep was applied in simulation, in which the motion of the motor link was controlled to be a sine wave with amplitude *A* and varying frequency *ω*:

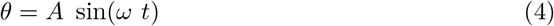

The frequency *ω* was varied from 1 to 10 rad s^−1^ in steps of 1 rad s^−1^ and for each frequency the amplitude *A* was manually tuned to reach the same angle deflection of the upper pendulum link *q*_1_ ≈ 1 rad. The applied amplitudes set to reach the deflection are summarized in Tab. 3. In every simulation the pendulum started from the rest position at 0 ° and the data was logged for 30 s with each frequency.

**Table 3.** Applied sine wave frequencies *ω* to the handle in a sweep scenario for the *P*_0_ and *P*_90_ pendulum to characterize the response of the systems. For each frequency, the sine amplitude *A* of the handle motion was manually tuned to reach a set deflection of the upper link (*q*_1_ ≈ 1 rad ≈ 60°) in the respective pendulum configuration.

| $\omega =$ | 1 | 2 | 3 | 4 | 5 | 6 | 7 | 8 | 9 | 10 | [rad s<sup>-1</sup>] |
| --- | --- | --- | --- | --- | --- | --- | --- | --- | --- | --- | --- |
| $A(P_0) =$ | 1.0 | 0.8 | 0.65 | 0.35 | 0.12 | 0.55 | 1.2 | 1.6 | 2.3 | 3.0 | [rad] |
| $A(P_{90}) =$ | 1.0 | 0.8 | 0.65 | 0.55 | 0.3 | 0.15 | 0.6 | 1.0 | 1.4 | 2.5 | [rad] |

Following the sweep, the two pendulum configurations were additionally excited with a sine signal where the amplitude *A* and frequency *ω* were randomly resampled every 100 time steps (=0.1 s). The amplitude was a pseudo-random value between 0 and 1, while the frequency was varied between 0 and 10 rad s^−1^. The random trial was repeated thrice per pendulum configuration, again starting from rest and logged for 60 s each.

The sweep experiment showed that the largest oscillation of the link side compared to the motor link motion was triggered at 5 rad s^−1^ and 6 rad s^−1^ for the *P0* and *P90* pendulum configuration, respectively (Fig. 12a). This was to be expected as the eigenfrequencies for the pendulum configurations were identified to be *f*_res(P0)_ = 0.78 Hz = 4.9 rad s^−1^ and *f*_res(P90)_ = 0.98 Hz = 6.16 rad s^−1^. At these frequencies the visualization in joint space also shows that the joint motions were closest to the expected nonlinear modes derived from the ideal conservative system. Additionally it could be observed that for both configurations, the phase shift between the motor link and the upper pendulum link moved from initially 0 for slow frequencies where the motor link and the pendulum moved synchronously towards an increasingly anti-phase relationship.

**Fig 12.**
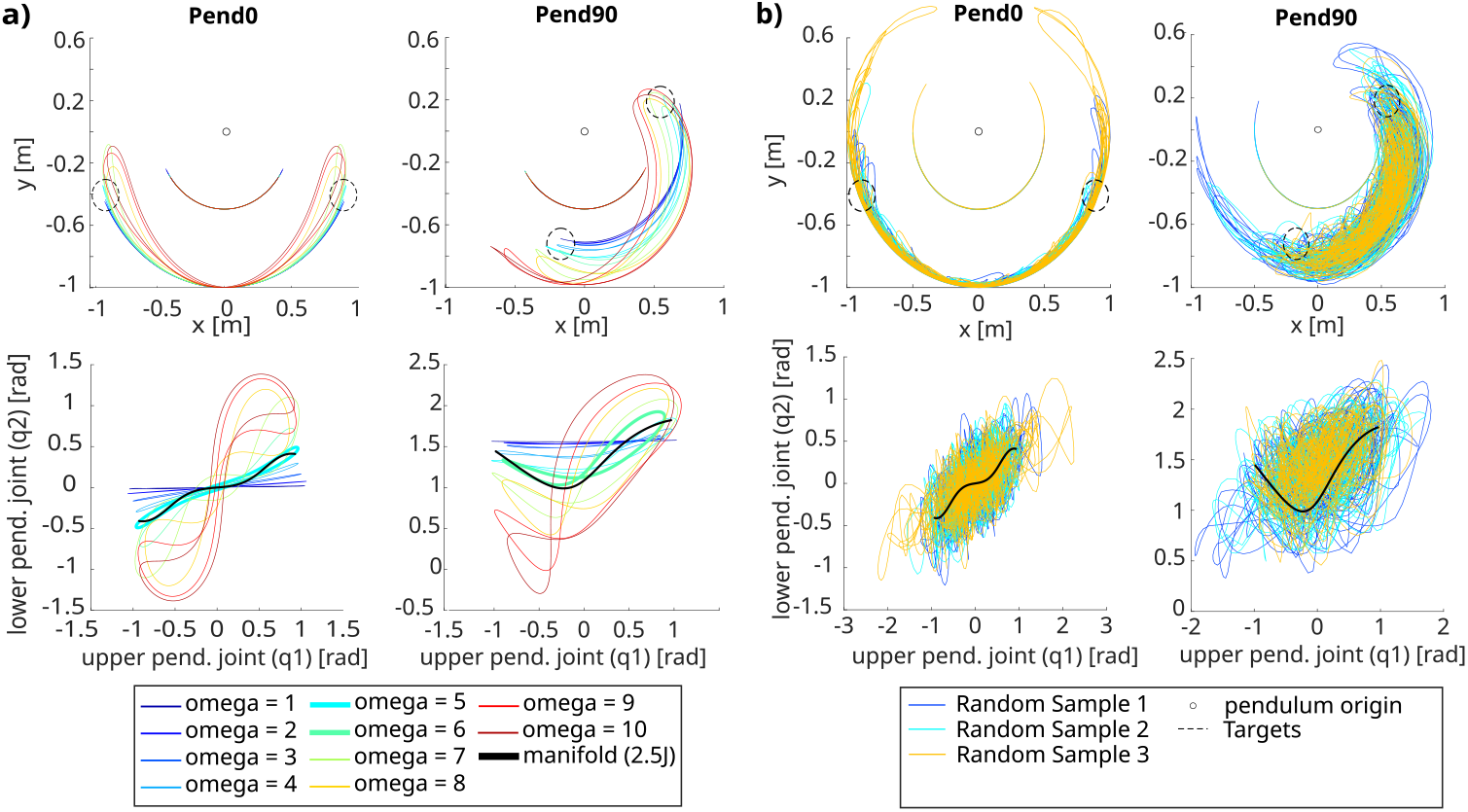
Comparison of the expected baseline strategies with the computed normal modes for the ideal system (solid black) and the averaged obtained data from the participants of the user study (dashed black). **a)** shows the comparison for the *P0* pendulum configuration in position (left) and velocity (right) space, while **b)** shows the respective plots for the *P90* configuration.

**Fig 13.**
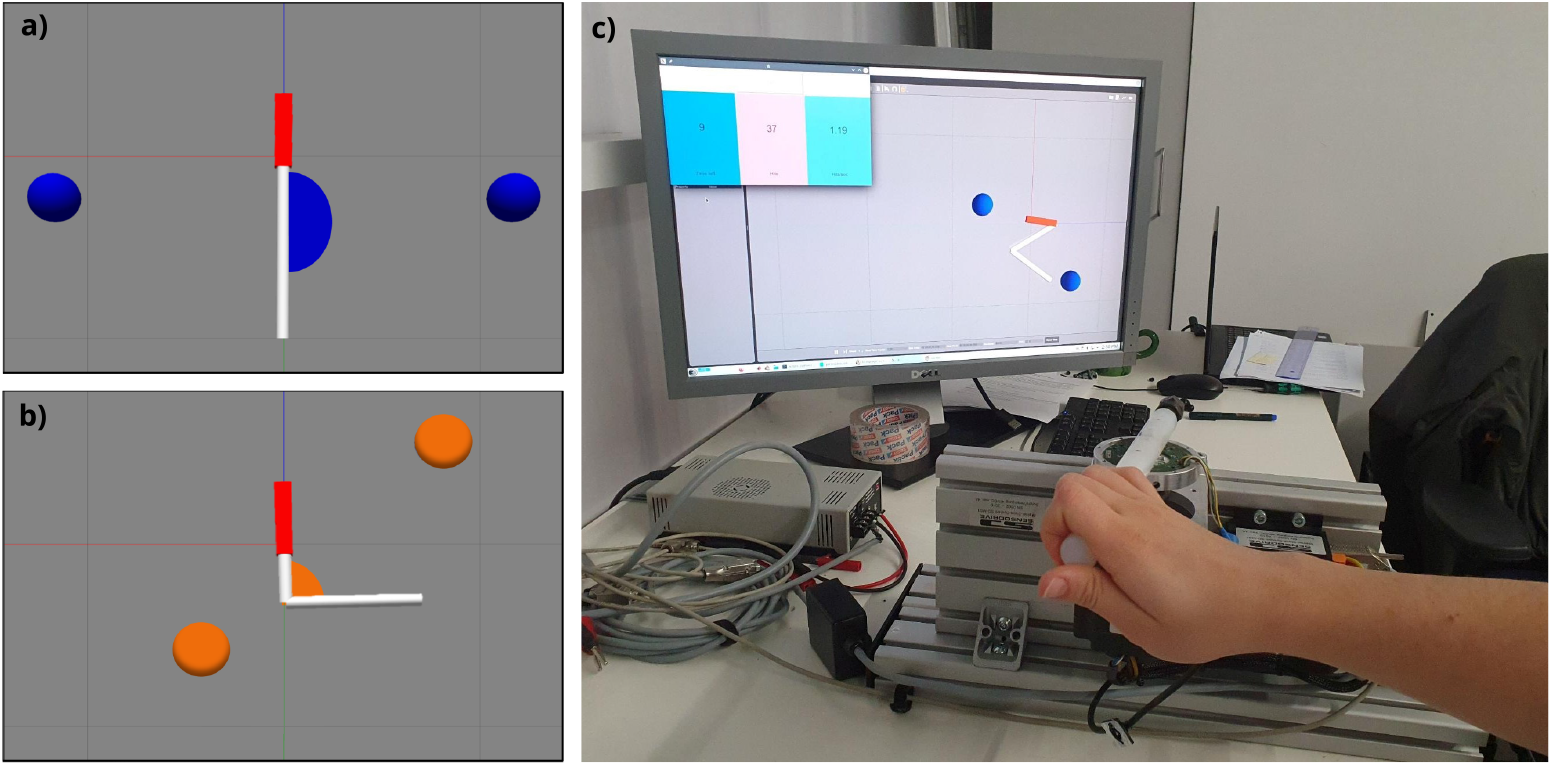
The targets for the designed high-score game are located such that the end point of the modes for the double pendulum configurations **a)** *P0* and **b)** *P90* lie within the radius of the targets. The radius of the target determines the required accuracy for the experiment task. **c)** During the experiment the the virtual double pendulum systems were visualized on-screen together with a GUI displaying the remaining time per trial and showed the participants the current hit count per trial.

The generation of the random motions on the motor link showed that the reachable space of the pendulum system is quite large, which is especially apparent in the joint space plots (Fig. 12b). Thus, the carried out characterization measurements prove that the pendulum systems can be excited in various ways and do not automatically fall to their respective eigenmode oscillation.

### Ethics statement

The experimental human user study was approved by the Institutional Review Board of the German Aerospace Center. The study included 20 right-handed participants (12 male, 8 female, between 21-45 years). Prior to the experiment, the procedure and objectives of the study were explained to all participants, and written informed consent was given to verify that they understood the written and oral task instructions.

### Experiment Task

Since the focus of this research is on dynamic interactions and it is known that humans apply very different strategies when moving slow [1, 46], the experimental task was defined to be a high-score game to encourage the participants to optimize the pendulum motions for speed and move rhythmically. The goal was to alternately hit two given targets as often as possible in a given time frame of 40 s. To test the stated hypothesis that humans indeed estimate and make use of the intrinsic motion patterns in compliant systems, the experiment task was defined such that it had an energetic benefit to exploit the pendulum system dynamics. Therefore, the targets were located so that the turning points of the systems’ nonlinear modes lied within the radius of the targets (Fig. 3c,d and Fig. 13a, b). The size of the radius determined the accuracy that the humans needed to achieve the task. A *hit* was considered when the lower link of the pendulum entered the target region, visually indicated by a color change of the targets. However, a point for the hit was only rewarded if the double pendulum did not swing through the target, i.e., leaving the target on the opposite side of entering to encourage the participants to keep a constant energy level. Whenever such an *overswing* occurred, next to the visual feedback, a beep sound was played as additional cue of error for the participant. Additionally, the lower link had to cross the zero-angle to initialize a new hit detection, such that the two targets on the mode end points needed to be hit alternately to collect points and thus exploit the whole motion range of the system modes. Earned points through correctly hitting one of the target balls and the remaining time per trial were displayed to the participants through a Python-based GUI (Fig. 13c).

### Experimental Variations

In order to investigate the robustness of the human control strategies to excite compliant systems in repetitive motions and scope how users adapt to different scenarios, different experiment variations were tested. The experiment task and instructions always remained identical, only system dynamics or target size and locations were altered. All Experiment variations were carried out within one session.

#### Experiment 1

The initial experiment setup focused on the interaction of the two main pendulum configurations *P0* and *P90* (Fig. 13a and b). For each pendulum configuration, the targets were arranged on the respective end points of the modes as previously described. Initially, the radius of the targets was set to be 0.1 m to allow some leeway for the participant in the required accuracy of the task. The pendulum parameters were set to the ones stated in Table 2.

#### Experiment 2

To test if and how the participants altered their control strategy for the *P0* and *P90* pendulum configurations in more challenging settings, the participants were divided into two groups. The reason to not test both experiment variations with all participants was to keep the experiment length within a reasonable time frame and thus avoid a bias in the results due to fatiguing effects.

For the first group, the settings in Experiment 2 were altered by changing the target size for the *P0* and *P90* pendulum configurations to 0.05 m, such that the task difficulty in terms of accuracy was higher. Therefore, the participants in this test group had to be more accurate to not overswing with the pendulum.

For the second group, the target size remained identical to the initial experiment run (0.1 m), but the mass of the lower link in the *P0* and *P90* configuration was changed to 0.625 kg. This mass alteration changed the behavior and dynamics of the pendulum systems, mainly noticeable to the participants by the slowed down oscillations of the system and increased feedback forces. Consequently, the *mode tool* was used again to re-compute the resulting modes for each pendulum configuration. However, it was decided to not change the positioning of the target balls in order to maintain an identical swing amplitude in Cartesian space and not bias the the human users in their control strategy. To still enable a fair comparison with the system mode, the considered energy level for the heavier system was lowered to 2.3 J, since then the mode end points still lied within the ball diameter of the targets that was used for the setup.

#### Experiment 3

To investigate the human control strategy to excite oscillating motions when the intrinsic system motions are not aligned with the task, i.e., the targets are not located on the end points of the computed nonlinear mode, a third pendulum configuration *P45* was investigated. In this configuration the spring offset between the upper and the lower pendulum link was set to be 45 °, thus representing an intermediate configuration between the two initial configurations (Fig. 9b). As for the two initial pendulum configurations, the parameters from Tab. 2 were used and the nonlinear mode of the conservative system was computed with the *mode tool* (Fig. 9a). However, for the *P45* configuration the targets were arranged to lie on a radius between the targets for the two previously described system configurations *P0* and *P90* (Fig. 9b and c), but still reachable with the excited pendulum.

The reason why a new pendulum configuration was chosen instead of relocating the targets for any of the previously described ones, was to avoid a bias in the control strategy of the users for these systems that could influence the overall behavior and make the interpretation of the results difficult. Instead, by choosing a slightly different variation of the same system provided with target locations that were systematically aligned with the targets of the *P0* and *P90* configuration, the three pendulum configurations could be presented and tested within one experiment run.

### Experiment Procedure

During the experiments, the participants were seated in front of a screen displaying the visualization of the pendulum simulations and the GUI showing the hit score and remaining time per trial. The haptic joystick was placed between the user and the display as shown in Figure 13c. The participants were free to orient themselves in front of the joystick handle and hold it in the way most comfortable to them. However, to minimize fatigue of the arm, the participants were asked to rest their elbow on the table.

Before the start of the experiment, each participant had a time span of 5 min to familiarize with the setup, the different configurations of the pendulum system and the task. For none of the experiment runs, were the participants instructed on how to excite the system to hit between the two targets, but the initial training time was granted to allow each participant to individually derive a strategy. After this time, the experimental trials were started. At the beginning of each trial, the Gazbeo GUI GzClient opened and displayed one of the investigated double pendulum configurations (*P0, P45, P90*), which were each presented four times sorted in a pseudo-randomized order individually generated per participant. Together with the simulation visualization, the GUI interface opened displaying the time countdown and the current hit score. It was also then, when the participant started to receive forces through the force-feedback joystick. The participant had then some seconds to tune into their control rhythm to oscillate the pendulum between the two targets. Only then a button in the python GUI was pressed by the experimenter to trigger the 40 s-countdown, initializing the counter of the hits and the recording of the data. In this way, solely the human strategy to sustain a system’s oscillation could be analyzed rather than the attempt of the participant to find their rhythm. After the countdown reached zero, all windows automatically closed and the reflected force of the joystick was set to zero, indicating the end of the trial to the participant. The recorded measurement data was then saved. Afterwards, the experimenter triggered the beginning of the next trial manually, to make sure the participant was ready. The trials were repeated until all pendulum configurations had appeared four times.

It was randomly selected whether a participant would start with the experiment run with the initial pendulum settings from Table 2, or with the respective alteration of the test group changing target size or lower link mass (Experiment 2). However, overall the experiment runs were arranged such that half the participants started with the initial settings and the other half with the respective alteration setup.

After one completed experiment run, the participants filled out a NASA TLX-inspired questionnaire about the perceived mental and physical work load.

The complete user study took around 60 min.

### Analysis and metrics

To analyze and compare the performance of different participants with regard to their intuitive ability to exploit the nonlinear normal modes of the double pendulum systems, the recorded data was post-processed with Matlab2020b. Each double-pendulum configuration (*P0, P45, P90*) was analyzed separately. For each configuration the worst of the four presented trials per person, judged by the achieved number of hit points, was identified and excluded from the data analysis. This was usually the first trial and was done to exclude initial learning effects or trials where the participant performed worse, e.g., due to a loss of focus or distraction. The considered time frame per trial for analysis included only the duration after the time countdown was started, i.e., the transient time each participant needed to find their rhythm was not analyzed. To equalize the recorded data, the periods were cut from the beginning and end of each trial, such that the data recording always started from one side and ended on the opposite. The three considered trials were then attached to one combined trial for analysis per pendulum configuration per person.

For each analysis trial, the turning points of the pendulum, i.e., where the target should be hit, were identified by finding the indices where the velocity of the pendulum tip reached zero in Cartesian space. The maximum Cartesian velocity was taken to identify the zero crossings, which was taken as cutting point between periods. Thus, each period included an attempt to hit the target on either side. Per period, it was identified whether the participant had hit each target through the recorded hit data, which indicated 1 for each *hit*, 0 if the target was not reached (*noreach*) and -1 for an *overswing* through the target. For comparison with the system modes, only the periods where the target of either side was reached were considered, i.e., the participant maintained the correct energy level. The data per participant per period were overlaid and then averaged to find the mean control strategy and performance per participant.

For each participant, descriptive information such as excited oscillation frequency of the different pendulum configurations, obtained hit points and number of periods in which the targets were not hit due to too little energy input or the hits were not counted due to overswinging, were identified. To investigate whether the participants applied a common control approach, it was also analyzed for each participants how the motor link, i.e., *θ*, was commanded in comparison the upper pendulum link coordinate *q*_1_. In this context, the phase lag between the human motion on the handle, i.e., motor link, and the upper pendulum link was identified and the maximum amplitude difference between the two coordinates per period was computed.

Additionally, the mode metric was introduced to quantify how close the excited pendulum motion through the participant was to the expected computed nonlinear normal modes of each pendulum configuration in the conservative case. The metric will be detailed in the following.

### Mode metric

Relating the angle positions and velocities of the upper link *q*_1_ and the lower link *q*_2_, the motions of the different pendulum configurations excited through each participant were mapped in joint space. There, they were compared with the the expected nonlinear normal mode of each system computed with the *mode tool* by the mode metric *η*.

This metric is based on the Dynamic Time Warping (DTW) principle to calculate the similarity of the excited pendulum motion of the humans and the computed normal modes of the ideal system taken as reference. The recorded pendulum signal from the experiment data is denoted *q*(*k*), while the reference coordinate is summarized in *Q*(*k*), where *k* is discrete time. Both *D*-dimensional signals are interpolated to have the same length *K* for comparison and the turning point at the same target size is taken as start point of the alignment. Since a nearest neighbor comparison would be prone to not include all features of the curves, and simply taking the euclidean distance between the interpolated points could lead to distortions due to slightly varying times per period per participant, DTW was chosen. Instead of only taking the Euclidean distance for the monotonously increasing points of both position vectors, DTW finds for each time instance *k* of *Q* a corresponding time index *f* (*k*) of *q* (*f* : N *→* N), such that

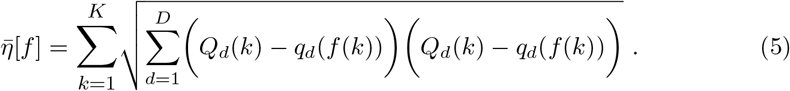

is minimized. The mode metric used for comparison is the minimum value of 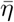 over all *f* found by DTW, i.e., 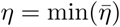. We used Matlab’s built-in `dtw`-function [47] for computation from the data. In a first approach the pendulum paths are solely compared in position space with *q* = [*q*_1_, *q*_2_] by *η*_pos_. However, to fully define a mode, the manifold including the velocities, 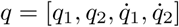, needs to be considered. Thus, additionally the DTW is applied to the state space of the experiment data to compare it to the respective trajectory on the manifold of the reference mode to calculate *η*_man_.

